# *Cutibacterium acnes* antibiotic production shapes niche competition in the human skin microbiome

**DOI:** 10.1101/594010

**Authors:** Jan Claesen, Jennifer B Spagnolo, Stephany Flores Ramos, Kenji L Kurita, Allyson L Byrd, Alexander A Aksenov, Alexey V Melnik, Weng R Wong, Shuo Wang, Ryan D Hernandez, Mohamed S Donia, Pieter C Dorrestein, Heidi H Kong, Julia A Segre, Roger G Linington, Michael A Fischbach, Katherine P Lemon

## Abstract

The composition of the skin microbiome varies widely among individuals sampled at the same body site. A key question is which molecular factors determine strain-level variability within sub-ecosystems of the skin. We used a genomics-guided approach to identify an antibacterial biosynthetic gene cluster in *Cutibacterium acnes* (formerly *Propionibacterium acnes*) that is widely distributed across individuals and skin sites. Experimental characterization of this cluster enabled the identification of a new thiopeptide antibiotic, cutimycin. Analysis of individual human skin hair follicles showed that cutimycin is an important factor regulating colonization resistance against *Staphylococcus* species.

**One Sentence Summary:** Cutimycin, a thiopeptide antibiotic produced by a widespread skin commensal, reduces *Staphylococcus* colonization of human follicles.

## Main Text

Niche competition among resident microbiota is postulated to influence skin microbial community composition via colonization resistance. In conjunction with host environmental factors (e.g., desiccation, low pH, high salt and high lipid concentrations (*1*)) and the host immune response (*2*), this results in a distinctive skin microbiota with variations among sites that are characterized as being predominantly sebaceous, moist or dry. Species of *Staphylococcus, Corynebacterium* and *Cutibacterium* (formerly the cutaneous *Propionibacterium* (*3*)) are among the most prevalent and abundant members of the human skin microbiota.

Recent studies have begun to uncover mechanisms of competition among bacterial species that shape microbiota composition across human skin (*4*). Small molecules are one means by which bacteria interact with each other and their environment. The genes required to produce these small molecules co-localize in biosynthetic gene clusters (BGCs) (*5*). BGCs are abundant in the human microbiome (*6*), but relatively few have a proven function (*6-8*). For example, the nasal and skin colonizer *Staphylococcus lugdunensis* produces a nonribosomal peptide, lugdunin, that inhibits growth of and colonization by *S. aureus* (*7*). Similarly, some strains of coagulase-negative *Staphylococcus* produce antimicrobial peptides that kill *S. aureus*. The levels of these strains are reduced in atopic dermatitis when *S. aureus* predominates and their expansion decreases skin colonization by *S. aureus* (*8*). Other mechanisms of bacterial competition on skin and in the nostrils include protease activity (*9*), disruption of *Staphylococcus* quorum sensing (*10, 11*), competition for iron (*12*), bacterial release of antimicrobial free fatty acids from host triacylglycerols (*13*), and niche competition mediated by the host (*14*). Nonetheless, the examples of lugdunin (*7*) and *Staphylococcus-*derived lanthipeptides (*8*) highlight the key roles of BGCs and their products for members of the human microbiota, and the need to identify and characterize secreted antibacterial compounds from other members of human skin and nasal microbiota (*15, 16*).

One widely distributed family of BGCs in the human microbiome is predicted to encode a sub-class of ribosomally synthesized, post-translationally modified peptides (RiPPs) known as thiopeptides (*6*). Many thiopeptides, such as berninamycin from *Streptomyces bernensis* and the semi-synthetic LFF571, which was used in a Phase 2 clinical study, have potent anti-staphylococcal activity via inhibition of protein synthesis (*17, 18*). We noted that the most widely distributed thiopeptide BGC in the human microbiome is found in 8/219 (3.7%) sequenced isolates of *Cutibacterium* species (Table S1), all *Cutibacterium acnes*. We and others predict that the product of this BGC (ppa0859–0866 in *C. acnes* strain KPA171202; Fig. 1A) is a thiopeptide (*19, 20*) structurally related to the antibiotic berninamycin (*21, 22*).

**Fig. 1.**
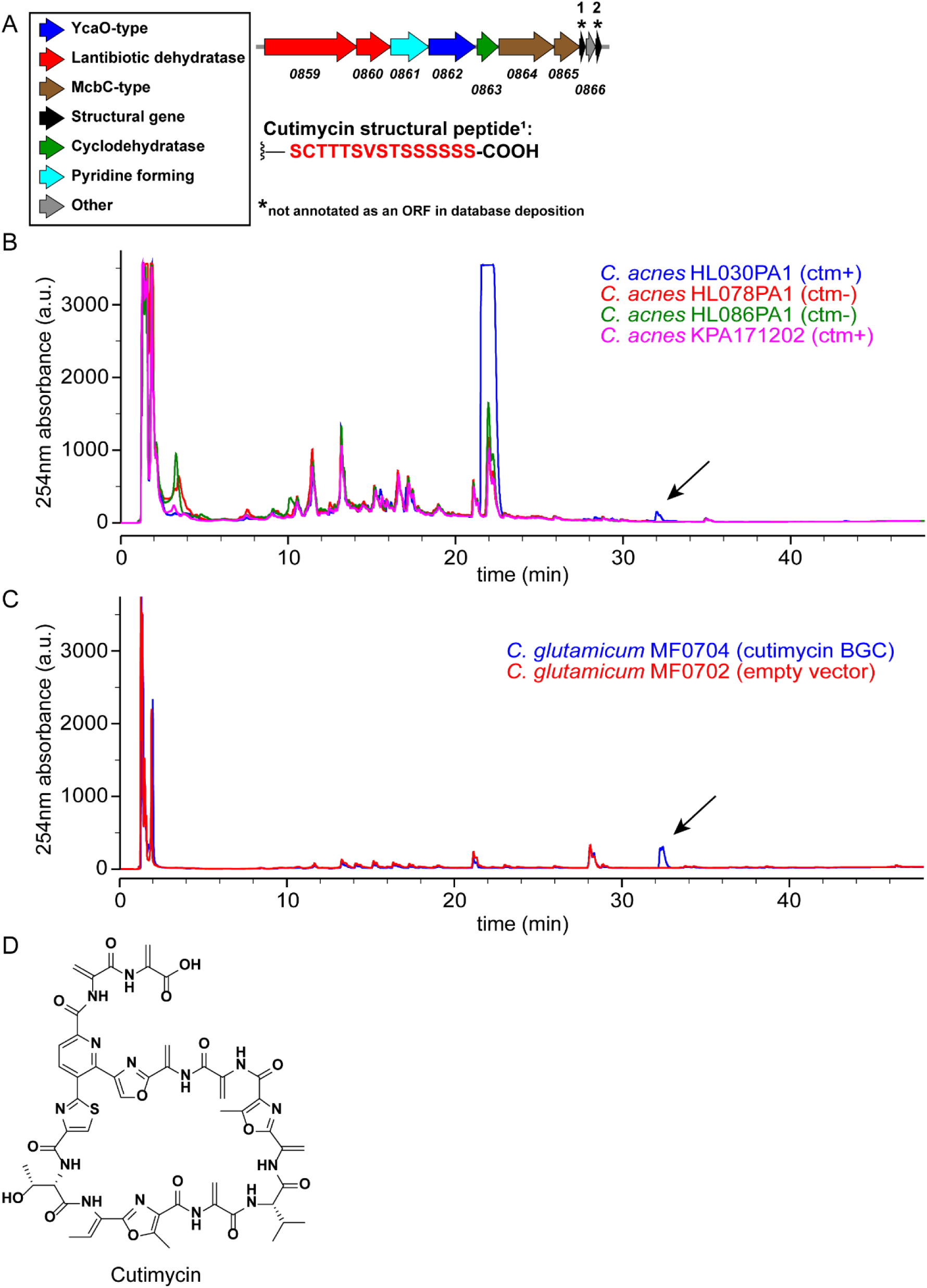
Detection of cutimycin (ctm), the product of BGC ppa0859-0866, in native and heterologous hosts. **A)** Arrow representation of the ppa0859-0866 BGC from *C. acnes* KPA171202. **B)** HPLC profiles for crude ethyl acetate extracts of select ppa0859-0866 BGC+ and - *C. acnes* strains. The thiopeptide product of ppa0859-0866, dubbed cutimycin, elutes at 73.5% MeCN as visible in the blue trace from *C. acnes* HL030PA1. Arrow indicates the cutimycin peak. **C)** Comparison of HPLC profiles of crude cell extracts of *C. glutamicum* hosting the cutimycin BGC on a plasmid (blue) versus hosting the empty vector control (red). Arrow indicates the cutimycin peak. **D)** Structure of the *Cutibacterium*-produced thiopeptide cutimycin.

*Cutibacterium* species are particularly well adapted to life on human skin with their ability to thrive in the lipid-rich environment of the human skin hair follicle, with its associated sebaceous gland. Hypothesizing that the small molecule product of this predicted cutimycin BGC plays a role in skin microbial community composition, we set out to identify this compound and determine its structure.

Based on its computationally predicted similarity to berninamycin (*23*), we hypothesized that the *Cutibacterium* BGC encodes a thiopeptide that *C. acnes* uses to target *Staphylococcus* species (phylum Firmicutes) in their shared habitats on human skin, including the skin inside the nostrils. To test for production of a thiopeptide, we selected phylogenetically distinct *C. acnes* strains that differ with respect to the presence or absence of BGCs. By analyzing crude culture extracts from these strains by HPLC, we observed that the thiopeptide BGC+ isolate *C. acnes* HL030PA1 produces a compound with a retention time and UV absorption spectrum similar to that of the thiopeptide berninamycin (Fig. 1B, blue trace).

To determine whether the BGC from *C. acnes* HL030PA1 was sufficient to produce the observed compound, we expressed this cluster heterologously in a related Actinobacterium, *Corynebacterium glutamicum*. We analyzed organic extracts prepared from cell pellets of the wild-type and BGC+ strains of *C. glutamicum*, observing that the latter but not the former contained a molecule identical to the one produced by *C. acnes* HL030PA1 (Fig. 1C). Unlike in the cell pellet, we did not detect the molecule in an extract of the culture supernatant of the BGC+ *C. glutamicum.* These data establish that the *Cutibacterium* BGC is sufficient for the biosynthesis of the molecule detected, but not its export.

Because of higher production yields and the ability to grow under aerobic conditions, we scaled up cultivation of the cutimycin BGC+ *C. glutamicum* and purified the thiopeptide (Fig. S1). Determination of the accurate mass at m/z 1131.3364 allowed prediction of a formula of C51H51N14O15S+, with a predicted monoisotopic mass of 1131.3373 (Δ_theoretical_ = 0.8 ppm) (Figs. S2 & S3). The planar structure of the thiopeptide was solved *de novo* based on 1D and 2D NMR experiments and HRMS^e^ and is described in depth in the supplemental information (Figs. S4-S11; Table S2). The configuration of each stereogenic center was determined using Marfey’s analysis (Fig. S12). We defined atom position numbering and the amino acid numbering (Figs. 1D and S13) following the convention set in the previous publication of the structure of berninamycin for ease of comparison (*24*). We assigned the trivial name cutimycin to this *Cutibacterium*-derived thiopeptide.

Cutimycin has potent activity *in vitro* against *Staphylococcu*s but not against commensal Actinobacteria from skin. Based on cutimycin’s structural similarity to berninamycin and LFF571 (Fig. S14), we hypothesized it would also display anti-staphylococcal activity but would lack activity against common Actinobacteria skin commensals. To test this, we determined the MICs for cutimycin and berninamycin against a selection of species commonly found on skin sites, including the nostrils (Table S3). Cutimycin exhibited potent inhibition of the USA300 community associated methicillin-resistant *S. aureus* strain NRS384 (MIC 0.2 μM), as well as strains of *Staphylococcus epidermidis*. In contrast, a panel of other *C. acnes* strains, with and without the cutimycin BGC, plus two common skin and two common nasal *Corynebacterium* species (phylum Actinobacteria) displayed increased resistance to cutimycin with MICs ≥ 3.2 μM. These data led us to hypothesize that cutimycin favors the growth of resident skin Actinobacteria, including *C. acnes*, over that of common skin staphylococcal species.

Cocultivation with susceptible *Staphylococcus* strains increased transcription of the cutimycin BGC. The production of secondary metabolites can be costly and BGCs often contain regulatory mechanisms for inducible rather than constitutive expression. We did not identify an obvious regulatory element in the cutimycin BGC. However, we hypothesized that cutimycin-susceptible species would induce transcription of cutimycin, whereas resistant species would not. To test this, we assayed transcription of the cutimycin BGC during *in vitro* cocultivation of *C. acnes* with *S. aureus, S. epidermidis* (both susceptible) or *Corynebacterium striatum* (resistant) compared to *C. acnes* monocultivation. For this qRT-PCR-based assay, we used *C. acnes* KPA171202 because the encoded cutimycin BGC (*25*) is transcribed during exponential growth *in vitro* (*19*). Compared to when *C. acnes* was grown alone, transcript levels of ppa0860 from the cutimycin BGC in *C. acnes* KPA171202 increased twofold in the presence of *S. aureus* or *S. epidermidis*, but decreased in presence of *C. striatum* (0.16 fold) (Fig. 2). These results indicate that cutimycin transcription is selectively increased by the presence of *Staphylococcus* targets. To determine whether cutimycin is produced *in vivo* in *C. acnes*’ natural habitat, we used mass spectrometry to analyze pooled content from hair follicles of intact skin of individual adult volunteers. Due to the small quantity of each sample (25-80 follicles), it was not possible to observe cutimycin using untargeted mass spectrometry (*26*). Using targeted mass spectrometry, we detected cutimycin in 28 % of the samples (Fig. S15, Table S4). Based on an external standard and a typical hair follicle volume of 0.2 mm^3^ (*27*), the concentration of cutimycin was estimated to be to be 0.97 +/-0.12 μM in the samples where the molecule was detected. However, the local concentrations likely exceed this value as cutimycin-producing strains are unlikely to be evenly distributed throughout the follicle. Based on these data, we hypothesized that cutimycin plays a role in modulating levels of *Staphylococcus* in the context of human skin colonization.

**Fig. 2.**
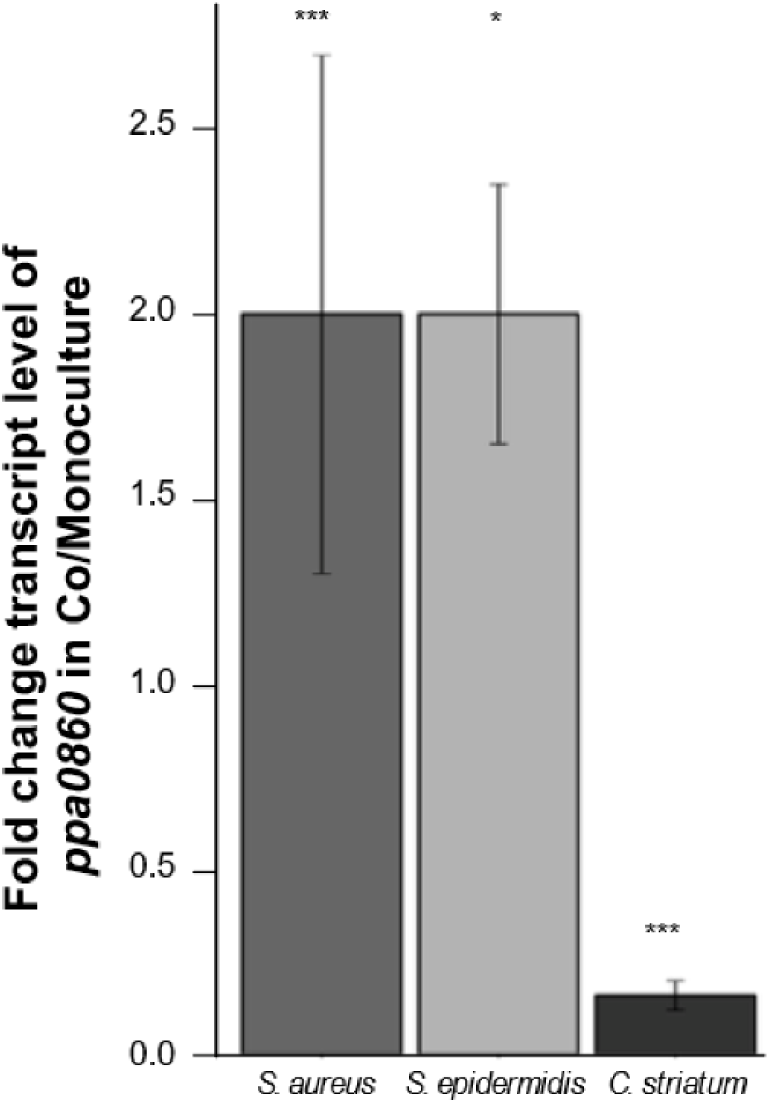
The *C. acnes* ppa0860 transcript increases during cocultivation with *Staphylococcus*. The ratio of qRT-PCR results for ppa0860 from co-vs. monoculture with *S. aureus* (*Sau*, n=8, *p*=0.0001), *S. epidermidis* (*Sep,* n=3, *p*=0.047) and *C. striatum* (*Cst,* n=3, *p*=0.00005). Paired t-test. Error bars are SD.

The cutimycin BGC is widely distributed on human skin but present in only a subset of strains at each body site. Humans harbor an abundance of *Cutibacterium* on their skin, with *C. acnes* being the overwhelmingly predominant species. In the absence of an animal model for *C. acnes* colonization of skin hair follicles, we began by examining isolate and metagenomic sequence data to explore a potential role for the cutimycin BGC in modulating the composition of the human skin community. The cutimycin BGC is present in the genomes of only about 4% (8/219 in Table S1) of sequenced *C. acnes* isolates. However, in a longitudinal high-resolution skin metagenomic dataset from 12 healthy volunteers (*28*), the cutimycin BGC could be detected in 11/12 individuals. When it was present in an individual, it could be identified across sampled sites stably over time (orange bars in Fig. 3A and S16). In spite of its broad distribution, in the majority of samples < 30% of the total *C. acnes* contained the BGC. *C. acnes* is also a common member of the human nostril microbiome, which is assayed from the skin of the nasal vestibule (*29, 30*). Because this site was absent in our metagenomic dataset, we also assayed for the cutimycin BGC in nostril metagenomic data from the Human Microbiome Project, finding it in 8/75 samples (10.7%). From the 12 intensively sampled volunteers, we also examined the distribution of four other *C. acnes* BGCs predicted to code for interesting bioactive compounds (Table S5). Similar to the cutimycin BGC, we observed a wide distribution with strain-level variation for these other *Cutibacterium* BGCs on adult human skin (Figs. 3A and S16). There were, however, no significant correlations between the absence/presence of the cutimycin BGC, or any other predicted *C. acnes* BGC, and the abundance of specific skin commensals in these 12 volunteers. At first these results seemed incongruous with cutimycin’s anti-staphylococcal activity. However, we reasoned that the spatial scale of a skin swab is much larger than the scale at which *C. acnes* interacts with neighboring bacterial species. Therefore, we hypothesized that a much finer spatial resolution was required to test whether cutimycin has an effect on microbial community composition. Because *C. acnes* is a known resident of human skin follicles, we next assayed the distribution of the cutimycin BGC and its potential impact on community composition at the level of individual human skin hair follicles.

**Fig 3.**
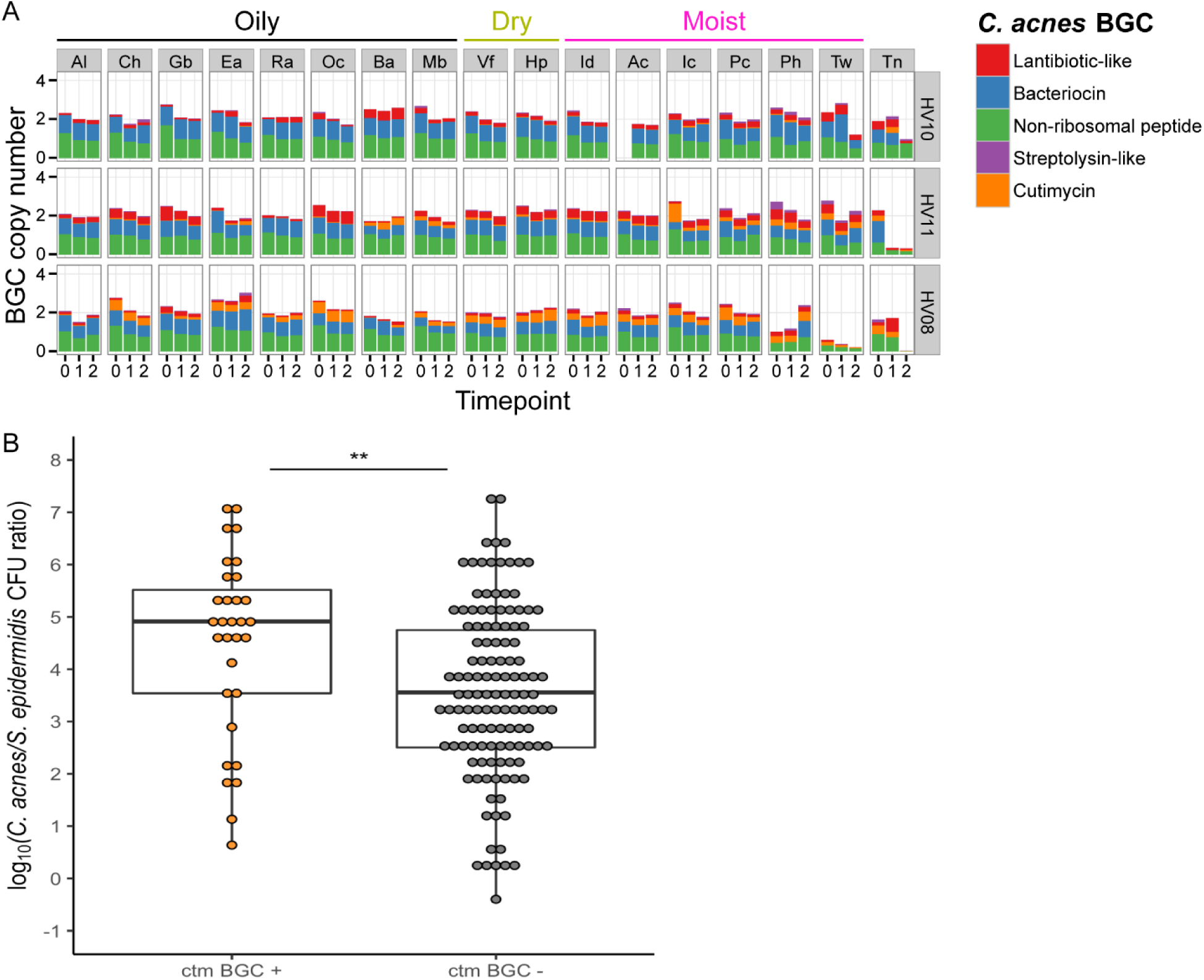
**A)** Example of spatial and temporal distribution of *C. acnes* BGCs in the skin metagenomes of three healthy individuals with low, medium or high relative abundance of the cutimycin BGC across 18 skin sites. Rows group samples from a single volunteer (coded Healthy Volunteer (HV) 01-12), columns represent samples across a specific body site (coded Ac through Vf). Cells contain bar graphs for each of the three time points, depicting the copy number of the *C. acnes* BGCs, which are standardized by comparing against 13 *C. acnes* housekeeping genes. The BGC types, bacteriocin, lantibiotic, non-ribosomal peptide, streptolysin-like peptide and cutimycin are color coded and their abundances are stacked on top of each other (a colored bar with a height of 1 means all of the *C. acnes* in the sample harbor that BGC, whereas a height of 0 means the BGC is absent in all of the *C. acnes*). Al = alar crease, Ch = cheek, Gb = glabella, Ea = external auditory canal, Ra = retroauricular crease, Oc = occiput, Ba = back, Mb = manubrium, Vf = volar forearm, Hp = hypothenar palm, Id = interdigital web space, Ac = antecubital fossa, Ic = inguinal crease, Pc = popliteal fossa, Ph = plantar heel, Tw = toe web space, Tn = toenail (see also diagram in Fig. S16A). **B)** Box plots of the impact of the cutimycin (ctm) BGC presence (+) or absence (-) on the log_10_ ratio of *C. acnes/S. epidermidis* CFUs from individual human skin follicular plugs (n=156, each follicle is represented by a dot) collected from 16 participants in total. For statistical analysis, data were pooled based on the assumption that follicles within an individual are independent. There was a statistically significant difference in Cac/Sep (Wilcoxon signed-rank test, *p*=0.006) between ctm+ and ctm-samples.

Skin follicles containing the cutimycin BGC have a higher ratio of *C. acnes* to *S. epidermidis*. To address the distribution question across individual skin hair follicles, we sampled the contents of 6-10 healthy follicles from the outer surface of the nose of 16 human volunteers (Fig. S17A, B and Table S6). For each follicle, we quantified colony forming units (CFUs) to assay community composition and performed PCR of *Cutibacterium* isolates to determine the presence or absence of the cutimycin BGC. From 156 individual follicles, we identified three bacterial species by 16S rRNA gene sequencing of representative colonies: *C. acnes, Cutibacterium granulosum*, and *S. epidermidis*. These three species were easily distinguished by colony morphology (Fig. S17C), permitting quantification of each in follicular content. Since follicles differed in the total number of CFUs (possibly due to size, sampling or true variation), we measured the ratio of *C. acnes*/*S. epidermidis* in follicles with and without the cutimycin BGC (Fig. S17D-E). Strikingly, follicles that contained *C. acnes* strains that were cutimycin BGC-positive had higher *C. acnes*/*S. epidermidis* ratio than those that were cutimycin BGC negative (Wilcoxon signed-rank test, *p=*0.006) (Fig. 3B). This finding suggests a role for cutimycin in modulating the composition of the skin microbiome, and it highlights that the characteristic spatial scale of this interaction is tiny – that of an individual follicle.

We also observed some cutimycin BGC-negative follicles with a high *C. acnes*/*S. epidermidis* ratio. One possible explanation is that alternative *C. acnes*-produced anti-*Staphylococcus* activities are present within these follicles, possibly encoded by one of the other predicted *C. acnes* BGCs (Fig. 3A and S16). However, other bacterial molecules/mechanisms could be mediating the competitive interactions, e.g., nutrient competition, toxic primary metabolites (*31*) or antimicrobial free fatty acids released from host triacylglycerols (*32*), as well as possible host-mediated effects.

In addition to inferences regarding the impact of cutimycin on community composition, these data on the presence or absence of the cutimycin BGC in individual human skin follicles indicate that follicles might be colonized by at least 2 different *C. acnes* strains and that *C. acnes* strain-level colonization is punctate, sometimes varying from follicle to follicle in an individual (Fig. S17D; Table S6). Thus, future exploration of the effect of *C. acnes* small molecules on human skin microbiota requires investigating at the fine-scale resolution of the individual skin follicle (Fig. S17C; Table S6).

Here, we have identified a molecular mechanism of niche competition between two of the most common members of the human skin microbiome: *C. acnes* and *Staphylococcus* species. With this work, cutimycin becomes one of the few BGCs from the human skin microbiome with a known molecular function. Our elucidation of cutimycin’s function will facilitate exploration into possible clinical applications of cutimycin to selectively inhibit *Staphylococcus* colonization, while leaving commensal Actinobacteria undisturbed. *S. aureus* nasal colonization is a risk for invasive infection (*33*) and, in the absence of an effective anti-staphylococcal vaccine (*34*), there is a need to identify strains of beneficial bacteria and their bioactive products that could be used to generate nasal and skin microbiota resistant to colonization by *S. aureus*. Such approaches might also have application in preventing or treating skin diseases that include a shift in either microbiome composition, such as in atopic dermatitis flares, or in host-microbe interactions, such as acne vulgaris (*35, 36*).

In conclusion, this successful elucidation of the cutimycin-mediated competition of *C. acnes* with *Staphylococcus in vivo* on humans demonstrates the power of combining systems-level approaches (e.g., *in silico* mining to characterize an antibiotic from the skin microbiome) with reductionist approaches (e.g., the *in vitro* cultivation of known bacterial colonizers of human skin follicles to explore of the impact of cutimycin at the fine-scale resolution of the individual skin follicle) to discover new mechanisms that underlie the ecology of the microbiome.

## Acknowledgments

We are deeply grateful to the participants who provided sebaceous samples from their skin hair follicles.

## Authors contributions

Conceptualization: JC, MAF, KPL. Investigation: JC, JBS, SFR, KLK, ALB, AAA, AVM, WRW, SW, RDH, MSD, RGL. Figures: JC, JBS, SFR, KLK, ALB, AAA, MAF. Writing of the original draft: JC, SFR, MAF, KPL. Writing, review and editing: JC, SFR, KLK, ALB, AAA, PCD, HHK, JAS, RGL, MAF, KPL.

## Data and materials availability

All data is available in the main text or the supplementary materials.

## Supplementary Materials

Materials and Methods

Figures S1-S18

Tables S1-S9

