## Supplementary Materials for "*Cutibacterium acnes* antibiotic production shapes niche competition in the human skin microbiome"

[1]Department of Cardiovascular and Metabolic Sciences, Lerner Research Institute, Cleveland Clinic, Cleveland, OH, USA [2]Department of Microbiology, The Forsyth Institute, Cambridge, MA, USA [3]Department of Oral Medicine, Infection & Immunity, Harvard School of Dental Medicine, Boston, Massachusetts, USA. [4]Department of Chemistry, Simon Fraser University, Burnaby, BC, Canada [5]Microbial Genomics Section, National Human Genome Research Institute, National Institutes of Health, Bethesda, MD, USA [6]Collaborative Mass Spectrometry Innovation Center, Skaggs School of Pharmacy and Pharmaceutical Sciences, and Center for Microbiome Innovation, University of California, San Diego, La Jolla, USA [7]Department of Bioengineering and Therapeutic Sciences, University of California, San Francisco, San Francisco, CA, USA [8]Department of Human Genetics, McGill University and Genome Quebec Innovation Center, Montreal, QC, Canada [9]Department of Chemical and Biological Engineering, Princeton University, Princeton, NJ, USA [10]Department of Molecular Biology, Princeton University, Princeton, NJ, USA [11]Dermatology Branch, National Institute of Arthritis and Musculoskeletal and Skin Diseases, National Institutes of Health, Bethesda, MD, USA [12]Department of Bioengineering and ChEM-H, Stanford University, Palo Alto, CA, USA [13]Division of Infectious Diseases, Boston Children's Hospital and Harvard Medical School, Boston, MA, USA

**This PDF file includes:**

Materials and Methods
Figs. S1 to S18
Tables S1 to S9

### Materials and Methods

#### Identification of the cutimycin BGC in sequenced *Cutibacterium* genomes:

A search for the ppa0859-0866 (aka cutimycin) BGC was performed on 219 currently available *Cutibacterium* genomes on NCBI as of 10/1/2018 (Table S1). An Align Sequence Nucleotide BLAST optimizing for more dissimilar sequences (discontinuous megablast) using the default parameters was performed on each of the genomes on 10/1/2018. The genomes were referenced through their accession number and aligned against the following fragment from the ppa0859 gene from *C. acnes* KPA171202:
ACCAGCAGGCTTACGGCACACCTGGAGTCTGTTATTTCAGTAACTGCGCCTCGTGAC
AGATCTCGACTAATCACATTAAAACGTGACGTGCATAACATGCGCGTGGAAAAAGT
GAAGCAAAGACTACTGCACTGTCAGGGCTGCACGGCTAGCAACATTATGGAGCCTG
TATCTAGTTGGATCGATACATGTCGATCATACGAAATACTTGAGCGTCGGTTGAGTC
AAGTATATGCGGACAGCCGAGACGAAGTGTCTCGTCACTACAGGTCCCTCTACCAG
GCTGACGCAATAGAGCCAAGCATTCAACTCTCAGGTGGCCAGCTCCACAGTGACC.

#### Strains and growth media:

All strains and plasmids used are listed in Table S7. *C. glutamicum* was grown aerobically using brain heart infusion (BHI). For other *Corynebacterium* species BHI was supplemented with 1% Tween 80 (BHIT). *Staphylococcus* species were grown aerobically in tryptic soy broth (TSB) or and *Cutibacterium* species were grown anaerobically using BHI broth. For Figure 2, all strains were grown on BHI agar.

#### Cloning of the cutimycin gene cluster:

All primers used in this study are listed in Table S8. The cutimycin gene cluster was cloned in SuperCosI (Agilent Technologies, Inc.) by a Gibson assembly strategy (*37*). The SuperCosI vector was linearized by PCR amplification using primers JC_Super_HL030_FWD/_REV and the cutimycin BGC was amplified in two parts from *C. acnes* HL030PA1 genomic DNA using primers JC_HL030_pt1_FWD/_REV and JC_HL030_pt2_FWD/_REV. The resulting 16 kb assembled construct, pJC215, was introduced into chemically competent *E. coli* Top10 cells and its entire nucleotide sequence was verified by DNA sequencing.

The cluster was subsequently subcloned using a two-step strategy into the *Corynebacterium–E. coli* shuttle vector pCGL0243 (*2*) for expression in *Corynebacterium*. First, pJC215 was digested with HindIII-HF (NEB) and the gel-purified 6376 bp fragment was introduced into HindIII-linearized pCGL0243. The resulting intermediate construct was linearized with BamHI to allow introduction of the second part of the cutimycin BGC, a 5.1 kb BamHI fragment from pJC215, yielding pJS004. The integrity of the construct and relative orientation of the inserts were verified by restriction digest and PCR (primers oKL244 and oKL424 to oKL439). The cutimycin expression construct (pJS004) and empty vector pCGL0243 were introduced into competent *C. glutamicum* DSM 20300 (ATCC 13032) as previously described (*38*). Transformants were selected on BHI + 20 µg/ml kanamycin.

#### Analytical detection of cutimycin production:

Cutimycin was detected in extracts from its native producer *C. acnes* HL030PA1 (Fig. 1B, blue trace) and the heterologous production host *Corynebacterium glutamicum* (Fig. 1C, blue trace). The latter was used for large-scale production because, 1) compared to *C. acnes*, it is easier to grow large quantities of *C. glutamicum* and 2) the occurrence of fewer co-eluting contaminants during further HPLC purification. A 50 ml BHI + kanamycin (25 µg/ml) preculture was inoculated from glycerol stock with the *C. glutamicum* MF0704 and grown in a baffled flask for 18 hours at 30°C, 225 rpm. A control culture was inoculated with the empty vector strain, *C. glutamicum* MF0702. (The producer preculture had a noticeable growth defect compared to the control.) An inoculum of 5 ml preculture was used per liter of production culture (in BHI + 25 µg/ml kanamycin). The production cultures were grown for 3 days at 30°C, 225 rpm. Production cultures were harvested by centrifugation at 8000 x g for 20 minutes, culture medium was discarded and the cell pellets combined and resuspended in $dH_2O$ (typically 25 ml $dH_2O$ was used per pellet from 1 liter culture). Cell clumps were homogenized by stirring (500 rpm on stir plate) and frozen in 45 ml aliquots (-20°C) until they were extracted.

For extraction, the 45 ml aliquots were completely thawed and $dH_2O$ was added to a total volume of 300 ml. This cell suspension was extracted 3 times with 300 ml ethyl acetate (EtOAc) in a separation funnel. Organic extracts were combined and dried with the rotovap. The dry residue was dissolved in 7 ml of MeOH and transferred to a smaller vial after filtering insolubles over a cotton plug. This solution was again rotovapped to dryness and then resuspended in 2 ml of MeOH for HPLC purification.

The MeOH-soluble part of the crude EtOAc extract was filtered over a cotton plug prior to HPLC analysis, using the Agilent SB-C18 column (1.8µm, 3.0x100mm). We used a gradient of $H_2O$ + 0.1% formic acid (FA) and acetonitrile (MeCN) + 0.1% FA, (98% $H_2O$, 2% MeCN for 3 min, ramped to 100% MeCN over 40 min and leaving constant at 100% MeCN for an additional 2 min) at a flow rate of 0.4 ml/min. Absorbance was monitored at 254 nm and cutimycin eluted at 32.3 min (or 73.5% MeCN).

**Purification of cutimycin:**

A dried extract of 0.458 g from *C. glutamicum* carrying the cutimycin BGC was subjected to a stepwise eluotropic gradient (50, 60, 70, 80, 90, 100% methanol with water, 40 mL fractions) on a 5 g Discovery $C_{18}$ prepacked column. The 80, 90, and 100 % methanol fractions were determined to contain cutimycin by UPLC-HR-ESI-TOFMS and were combined and concentrated to dryness *in vacuo* to give a yellow foamy oil. The product was resuspended in 2 mL of methanol and the resulting solution purified on an Agilent 1100 series HPLC equipped with a quaternary pump, a DAD detector and a 1260 Infinity series fraction collector using a Phenomonex Kinetix 5 µm XB-C18 150 x 4.6 mm column. A gradient of MeCN + 0.02% formic acid, MeOH + 0.02% formic acid, $H_2O$ + 0.02% formic acid (45 % MeOH, 30% MeCN, 25% $H_2O$ for 2 min, ramped to 54% MeOH, 30%MeCN, and 16% $H_2O$ over 9 min) at a flow rate of 1.5 mL $min^{-1}$ was used for the purification. After each gradient the column was washed with 90% MeOH/10% MeCN for 5 min and equilibrated at the initial conditions for 5 min. The solvent mix was necessary to avoid peak fronting and tailing observed when using either methanol or acetonitrile alone. The peak at 9.3 min (*m/z* 1131.2 $[M+H]^+$) was collected (Fig. S1) and dried under vacuum yielding 1.8 mg of pure material as a white powder.

**Structure elucidation of cutimycin:**

Standard 1D and 2D NMR experiments ($^1H$, $^{13}C$, COSY, TOCSY, ROESY, HSQC, and HMBC) were acquired in DMSO-d6 in a 5 mm Shigemi tube at 600 MHz (151 MHz for $^{13}C$). Initial analysis of the $^1H$ (Fig. S4) and $^{13}C$ (Fig. S5) spectra indicated the presence of a complex peptide containing a large number of aromatic residues. Starting from the C-terminus at position

2 (Fig. S12), two consecutive dehydroalanine moieties could be assigned based on chemical shifts, the presence of the terminal alkenes, and HMBC correlations along the amide backbone (Fig. S11). The terminal carboxylic acid was determined by the observation of the $b_{15}$ and $b_{14}$ fragments observed in the high energy $MS^e$ scan (Fig. S3) with neutral losses of 18.0111 *m/z* (calcd 18.0106) and 87.0333 *m/z* (calcd 87.0320). This tail could be connected to the core of the molecule by HMBC correlations from both the amide proton at position 9 ($\delta_H$ 10.56) and the aromatic proton at position 12 ($\delta_H$ 8.26) to the carbonyl carbon at position 10 ($\delta_C$ 161.50) (Fig. S8-10).

The presence of two aromatic proton doublets at $\delta_H$ 8.26 and 8.53 that showed HMBC correlations to three quaternary carbons ($\delta_C$ 130.23, 146.85, and 149.40) unique to that spin system was indicative of a tri-substituted pyridine ring, a well-known motif in highly aromatized bacterial cyclic peptides. The relative positions of the aromatic protons were assigned based on the downfield shift of the carbon at position three on the pyridine ring, coupled with the strong three-bond HMBC correlation from the proton at $\delta_H$ 8.26 to the carbonyl at position 10 ($\delta_C$ 161.50). The substitution pattern of carbons 15 and 14 and relative connectivity to carbons 16 and 61 were assigned based on a strong three-bond HMBC correlations from the proton at position 13 ($\delta_H$ 8.53) to quaternary carbon 61 at $\delta_C$ 163.03 and a weak four-bond HMBC correlation from position 13 to quaternary carbon 16 at $\delta_C$ 139. 13 (Figs. S8 and S10).

An oxazole moiety was located at carbon 15 on the pyridine ring, based on the presence of a protonated methine (position 17, $\delta_H$ 8.75, $\delta_C$ 140.59) that exhibited strong HMBC correlations to two aromatic quaternary carbons (position 16, $\delta_C$ 139.13 and position 18, $\delta_C$ 158.06) with carbon chemical shift values consistent with the oxazole functional group. The next two amino acids in the chain were both identified as dehydroalanine based on the presence of diastereotopic terminal alkene protons (position 20, $\delta_H$ 5.74, 5.75 and position 24 $\delta_H$ 5.73, 6.35) with HMBC correlations to carbon atoms 19 and 23 respectively (Fig S9). NMR data alone was insufficient to extend this subunit, however a number of important additional HMBC correlations were observed, including correlations from the amide proton at position 25 ($\delta_H$ 9.37) and a methyl singlet at position 28 ($\delta_H$ 2.62, $\delta_C$ 11.45) to the same quaternary carbon (position 26, $\delta_C$ 159.49). Methyl 28 also displayed HMBC correlations to two other quaternary carbons ($\delta_C$ 154.21 and $\delta_C$ 129.17) suggestive of a highly substituted aromatic system at the eastern terminus of subunit A (Fig. S11A).

To complete the southern component of subunit A, a similar strategy was employed. The substituent at carbon 14 of the pyridine ring was defined as a thiazole ring based on the presence of a protonated singlet methine (position 60, $\delta_H$ 8.43, $\delta_C$ 126.68) that showed HMBC correlations to quaternary carbons at $\delta_C$ 163.03, 159.83 and 149.40 (positions 61, 58 and 59). These signals could be distinguished as the carbonyl at position 58 and the quaternary carbon at position 59 based on their respective chemical shifts and the HMBC correlation from the amide proton at position 57 ($\delta_H$ 7.99) to the carbon at position 58 ($\delta_C$ 159.83). The next two amino acids in the chain were identified as L-threonine and dehydrobutyrine based on $^1$H, COSY and HMBC data. Finally, all three of the dehydrobutyrine proton signals possessed HMBC correlations to carbon 48 ($\delta_C$ 156.65), from which point no further structural information was available from the 2D-NMR data.

A contiguous chain of amino acids comprising dehydroalanine, L-valine, and dehydroalanine was identified in subunit B (Fig. S11B) from $^1$H, COSY and HMBC data. At the N-terminus of this chain (position 43) the amide proton and an adjacent methyl group (position

47 $\delta_H$ 2.59, $\delta_C$ 11.59) showed HMBC correlations to a carbonyl carbon at position 44 ($\delta_C$ 159.60). Additionally, methyl singlet 47 possessed HMBC correlations to two other quaternary carbons ($\delta_C$ 153.56 and $\delta_C$ 133.36). The downfield shift of the methyl group and the chemical shifts of these quaternary carbons suggested the presence of an aromatic heterocycle. Based on the remaining atoms from the molecular formula ($C_{10}N_2O_4$) and the unresolved HMBC correlations from the termini of both subunits A and B the presence of a pair of 4-carboxy-5-methyloxazole moieties was proposed. This hypothesis was well supported by the distributions and chemical shifts of the unresolved HMBC correlations from subunits A and B. The orientations of these two functional groups were assigned based on the HMBC correlations from methyls 29 and 47 to the carbonyl carbons at 26 and 44 respectively. Finally, the complete planar structure of cutimycin was completed by closing the peptide macrocycle via the Dha-Val-Dha segment between the newly installed methyloxazole subunits (Fig. S11B).

**Mass spectrometry of the cutimycin molecule:**

The sequence of the peptide was confirmed by UPLC-$MS^e$ analysis using a Waters i-Class Acquity UPLC coupled to a Waters Synapt-G2Si mass spectrometer operated in resolution $MS^e$ mode providing low energy (parent) and high energy (fragment) data at a resolution of 20,000 (Figs. S2 & S3). The mass spectrometer was calibrated using sodium-formate and the accurate mass was corrected using two lock masses from leucine encephalin. The separation was performed using a HSS T3 1.8 µm 2.1 x100 mm column at a constant flow rate of 0.5 mL/min using a gradient of 5 to 100% MeCN in $H_2O$ modified with 0.1% formic acid with a curve profile of 6 after a 0.3 min hold at the initial conditions. The curve profile was set to 6 yielding a linear gradient. Raw MS data was imported into Waters Unifi software and the correct peak annotated by importing the predicted structure as a scientific library and using a general accurate mass screening method to annotate molecular fragments in the $MS^e$ (high energy) spectra (Figs. S3 and S18; Table S9).

**Marfey's Analysis:**

The configurations of the two chiral amino acids were determined using Marfey's analysis (Fig. S12). 100 µg of cutimycin was heated with stirring at 95°C in 1 mL of 6 N HCl overnight, cooled, and the solution concentrated to dryness under a stream of air. To perform the derivatization 0.5 mL of 1% N$\alpha$-(2,4-dinitro-5-fluorophenyl)-L-alaninamide (Sigma) in acetone, 0.1 mL of 1M $NaHCO_3$, 0.05 mL of DMSO, and 0.1 mL of $H_2O$ were added to the vial and the resulting yellow solution stirred at 40°C for 1 hr. The same proportions at a tenth of the scale were used to derivatize 0.1 µmoles of each of the six amino acids L-valine, D-valine, L-threonine, D-threonine, L-*allo*-threonine, and D-*allo*-threonine. After cooling, each solution was evaporated to dryness and resuspended in 1.0 mL of 50% MeOH in $H_2O$.

Marfey's products were analyzed with a Waters Acquity i-Class UPLC coupled to a Synapt G2Si mass spectrometer operated in resolution mode. The derivatives were separated on an Acquity UPLC HSS T3 1.8 µm 2.1 x100 mm column at a constant flow rate of 0.5 mL/min using a gradient of 5 to 100% MeCN in $H_2O$ modified with 0.1% formic acid with a curve profile of 7 after a 0.3 min hold at the initial conditions. The gradient curve was set to 7 to give the gradient a convex shape. The first 0.6 min of eluent was sent to waste to avoid sending excessive salts into the source of the instrument.

**MIC determination:**

We assayed for the inhibitory activity of cutimycin against a set of skin bacteria, including *C. acnes* strains (KPA171202, HL030P A1, HL110PA1, HL086PA1, HL053PA2), *Staphylococcus* strains (*aureus* USA300, NRS384 and UAMS-1 and *epidermidis* W23144, DSM20042 and ATCC 35984) and *Corynebacterium* strains (*accolens* ATCC 49725, *jeikeium* DSM7171, *striatum* DSM20668 and *pseudodiphtheriticum* DSM44287) (Table S3). These data were compared to the activity of the structurally related compound berninamycin under the same circumstances. *Corynebacterium* spp. and *Staphylococcus* spp. were grown aerobically at 37°C overnight in BHIT or TSB, respectively, and inoculated into their respective fresh assay medium at a final dilution of 1/2500. *Cutibacterium* spp. were grown anaerobically at 37°C in BHI for four days before inoculating the assay medium at a 1/25 final dilution. Cutimycin and berninamycin were added to the assay medium at final concentrations of 0, 0.05, 0.2, 0.8 and 3.2 µM, and MICs were determined after incubation for 16 hours for *Corynebacterium* spp. and *Staphylococcus* spp. and three days for *C. acnes*. The obtained MICs were generally consistent between biological duplicates, but when a difference was observed, the highest value is reported.

**Transcriptional analysis of the cutimycin BGC in co- versus monoculture:**

*C. acnes* KPA171202 was grown for two days prior to addition of a second species to compensate for differential growth rates. First, *P. acnes* KPA171202 was resuspended to an $OD_{600}$ of 0.1 in sterile BHI medium and then sixteen 5 microliter spots were inoculated onto a 0.2-micron sterile polycarbonate membrane forming a grid pattern on BHI agar medium. The cultures were incubated under anaerobic conditions at 37°C and subsequently transferred to a fresh agar medium after two days of growth. To perform the co-culture assay, *S. aureus, S. epidermis* or *C. striatum* were each individually inoculated at an $OD_{600}$ of 0.1 onto the top of *C. acnes* spots. After an additional 20 hours of growth under anaerobic condition at 37°C, cells were harvested by immediately transferring cells plus membranes to TE buffer (10 mM Tris-HCl pH 8, 1 mM EDTA).

For RNA extraction, cells were resuspended in TE then incubated 15 minutes at 37°C with lysozyme (Ready-LyseTM; Epicentre) prior to RNA extraction using the Masterpure kit (Epicentre) coupled with 4 bead beating cycles (MPbio - FastPrep-24; 6 M/s 30 seconds) in 2 ml Lysing Matrix B bead tubes (MP Biomedicals). The extracted nucleic acids were treated with DNase for one hour using TURBO DNA-free (Ambion) according to the manufacturer's protocol. RNA was resuspended in 40 µl of nuclease-free dd$H_2O$. RNA quality was analyzed with an Agilent bioanalyzer using a total RNA pico chip.

For quantitative-RT-PCR, we used the qScriptTM One-Step SYBR® Green qRT-PCR Kit (Quanta Biosciences) according to the manufacturer's protocol. cDNA synthesis and PCR amplification were carried out in the same tube (96 wells plate Roche), followed by 10 minutes of reverse transcription at 50°C in a LightCycler® LC 480 (Roche) and then 40 cycles of real time PCR. The relative expressions were calculated using the Ct value determined through the Roche program. A student T-test was used for the statistics

**Targeted mass spectrometry of follicular content:**

A pool of follicular plugs per participant were assessed through mass spectrometry. Two independent experiments have been carried over using different extraction protocols. For the first experiment, samples came from a few volunteers and were identified by subject. For the second experiment, the content of ~25-80 follicles per subject were pooled for 5 volunteers.

In the first study, Eppendorf microtubes with pooled sample (5-20 follicular plugs) were removed from -80 °C freezer and warmed at room temperature for 15 minutes, 20 µL of acetonitrile added, sonicated for 10 minutes and allowed to extract for 72 hours at 4 °C. The supernatant from each sample was then transferred into a separate Eppendorf, the samples were lyophilized and reconstituted in 20µL of methanol with sonication.

For the second study, the Eppendorf microtubes with pooled sample (25-80 follicular plugs) were removed from -80 °C freezer, warmed at room temperature for 15 minutes and dissolved in 100 µL of ethyl acetate. The samples were sonicated for 10 minutes, centrifuged at 15,000 rpm for 5 minutes, and 80 µL aliquot was transferred into a separate Eppendorf for each individual sample. The same procedure was repeated 3 times, and three transferred aliquots were combined for individual samples. The ethyl acetate supernatants were then lyophilized and 50 µL, aliquot of methanol was added to re-dissolve the dry residues and sonicated for 5 minutes. A 110 µL of HPLC-grade methanol was added to the dry residue of hair follicles, the samples sonicated for 10 minutes, allowed to extract for 4 hours and centrifuged at 15,000 rpm for 10 minutes. The 100 µL aliquot of the supernatant was transferred into the Eppendorf to combine with the methanol-reconstituted dry ethyl acetate extraction residue. The samples were centrifuged at 15,000 rpm and 100 µL of each sample was transferred onto a 96-well mass spectrometry plate.

Thermo TSQ Access Quantum MAX QqQ instrument with HESI-II probe source (Thermo Fisher Scientific, Waltham, MA USA) was employed for targeted MS analysis. The following probe settings were used: spray voltage of 3500 V, sheath gas (nitrogen) pressure of 35 psi, auxiliary gas pressure of 10 psi, ion source temperature of 270 °C and auxiliary gas heater temperature at 440 °C. Spectra were acquired in single ion monitoring (SIM) and multiple reaction monitoring (MRM) modes. Data acquisition parameters were set as follows: minutes 0–0.5 were sent to waste; minutes 0.1–12 were recorded with collision gas pressure of 1.5 mTorr, isolation width of m/z 0.2 and with time of 0.05 s for each transition ion.

Direct infusion of cutimycin standard solution was used to optimize collision energies on Thermo TSQ Access Quantum MAX QqQ and ensure optimal fragmentation conditions for SIM ad MRM measurements. Calibration curve for cutimycin was built by injecting 10 µL of standard solution in methanol for series of dilutions from 1 µM to 10 nM concentrations for both instruments; 3 technical replicates for each concentration point. The most abundant ions in the CID spectrum were determined and correspondingly the Thermo TSQ Access Quantum MAX QqQ was set to perform two SIM scans for cutimycin +1 (m/z 1131.34) and +2 (m/z 566.17) ions as well as 2 MRM transitions of two most abundant ions: m/z 566 -> 912 and m/z 566 -> 632.

For the detection of cutimycin in follicular content, 10 µL of the prepared samples were injected and chromatographically separated using a Vanquish UPLC (Thermo Fisher Scientific, Waltham, MA), on a 100 mm × 2.1 mm Kinetex 1.7 µM, C18, 100 Å chromatography column (Phenomenex, Torrance, CA), 40 °C column temperature, 0.5 mL/min flow rate, mobile phase A 99.9% water (J.T. Baker, LC–MS grade), 0.1% formic acid (Thermo Fisher Scientific, Optima LC/MS), mobile phase B 99.9% acetonitrile (J.T. Baker, LC–MS grade), 0.1% formic acid (Fisher Scientific, Optima LC–MS). The solvent gradient was as follows: 0–1 min 2% B, 1–5 min ramp to 60% B, 5-8 min ramp to 100% B, 8-9.9 hold at 100% B, 9.9-10 min, ramp down to 2%B, 10–12 min equilibration at 2% B.

The obtained data were examined using Thermo Scientific Xcalibur software. The presence of a peak in both SIM and MRM spectra at the retention time that corresponds to the elution of cutimycin standard was considered an evidence of cutimycin detection (Figure S15).

Cutimycin was detected in 30% of all samples in the first study, and 20% in the second, with 28% rate of detection overall across both studies (Table S4). The range of detected amounts of cutimycin was 0.42 pmol to 0.60 pmol (47.80 to 67.76 ng) per hair follicle. This is the lower boundary amount, as most likely not all of the compound is recovered by either of the employed extraction procedures. For a typical volume of follicle of 0.2 cubic millimeters, these detected amounts correspond to concentration of cutimycin in hair follicles ranging from 0.84 to 1.19 µM (average of 0.97 +/- 0.12 µM). However, the local concentrations across follicles may vary and exceed this value.

**Metagenomic *Cutibacterium* BGC analysis:**

The distribution of *C. acnes* BGCs was analyzed for in skin metagenome for 12 healthy individuals, spanning 18 body sites and at 3 different time points (*28*). For this, a non-redundant list of *C. acnes* genes was created from all sequenced isolates in the databases, yielding a pan-genome of 3774 unique genes (*28*). Bowtie2 was used to identify what genes from the pan-genome are present in each sample from the metagenomics dataset. A gene was counted as present if over 40% of its length was covered by the sequence reads. This criteria reduces gene calling due to spuriously mapped reads or reads from orthologs of closely related species (*39*). Average coverage of each gene was calculated with samtools (*40*) and then normalized by the average coverage of 13 single copy marker genes (*41*) to yield a copy number estimate. We identified the presence of *C. acnes* BGCs by associating genes in the pangenome to those in Table S5.

The presence of the cutimycin BGC cluster is represented in four categories depending on how many genes could be detected with confidence among the reads within a sample. The categories are “none” (no genes from that BGC detected), “less than half”, “greater than half” and “complete” (all genes from the BGC detected in the sample). Copy number of a particular BGC was calculated based on the average copy number of all genes within the individual cluster.

**Analysis of the content of individual human follicles:**

All participants provided informed consent and samples were collected under a protocol approved by the Forsyth Institutional Review Board. Follicular content was harvested from pilosebaceous units on the outer surface of the human nose using Bioré Deep Cleansing Pore Strips (Kao Brands Company, Cincinnati, OH) similarly to a previously described procedure (*36*). We immediately picked six to ten individual follicular plugs from each participant’s fresh pore strip with a sterile P2 pipette tip and resuspended each separately in 100 µl BHI. Serial dilutions of up to $10^{-4}$ were performed and plated on BHI agar. After 4 to 5 days of growth under anaerobic conditions, we calculated the CFUs for the three visible colony morphologies, which corresponded to *C. acnes*, *C. granulosum* and *S. epidermidis*. Species assignment for each colony morphology was based on 16S rRNA gene sequencing of a representative set of colonies. For 11 participants, six to ten single colonies from each follicular plug with the typical *C. acnes* morphologically were colony purified and saved as frozen stocks in a 96-well format. To assay for the presence of the ppa0859 gene (i.e., the cutimycin BGC), a 15 µl colony PCR was performed on each of the 1208 bacterial isolates (or on a pool, Table 6) using the 2X Gotaq Green Master Mix (Promega) and 10µM of the oKL535 and oKL536 primers. We used *C. acnes* strains KPA171202 and HL086PA1 as positive and negative controls, respectively, for presence and absence of the ppa0859 gene in the PCR screen for the cutimycin BGC. To genotype the isolates as either ppa0859 positive or negative, each amplicon was subjected to electrophoresis on a 1% TAE agarose gel. The CFU ratio of *C. acnes* to *S. epidermidis* was calculated and corrected by adding one to the CFUs of *S. epidermidis* to account for when its value was zero. A Wilcoxon signed-rank test was performed on the ratio to compare the cutimycin positive and the cutimycin negative follicles.

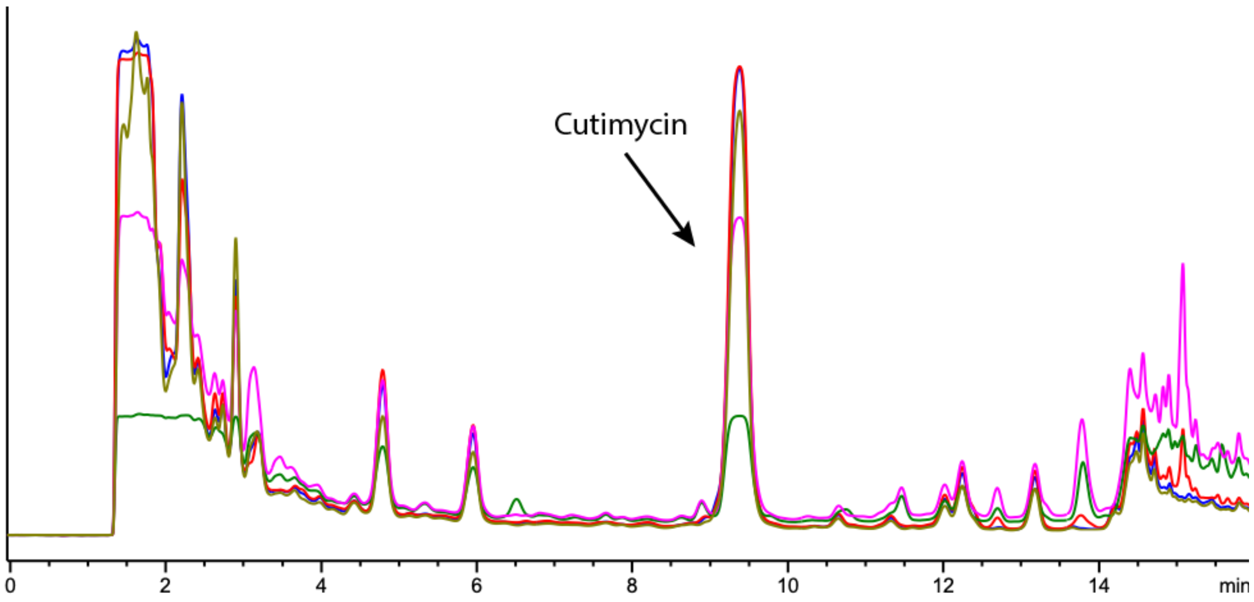


**Fig. S1**. HPLC chromatogram of cutimycin purification. The peak eluting at 9.3 minutes was collected.

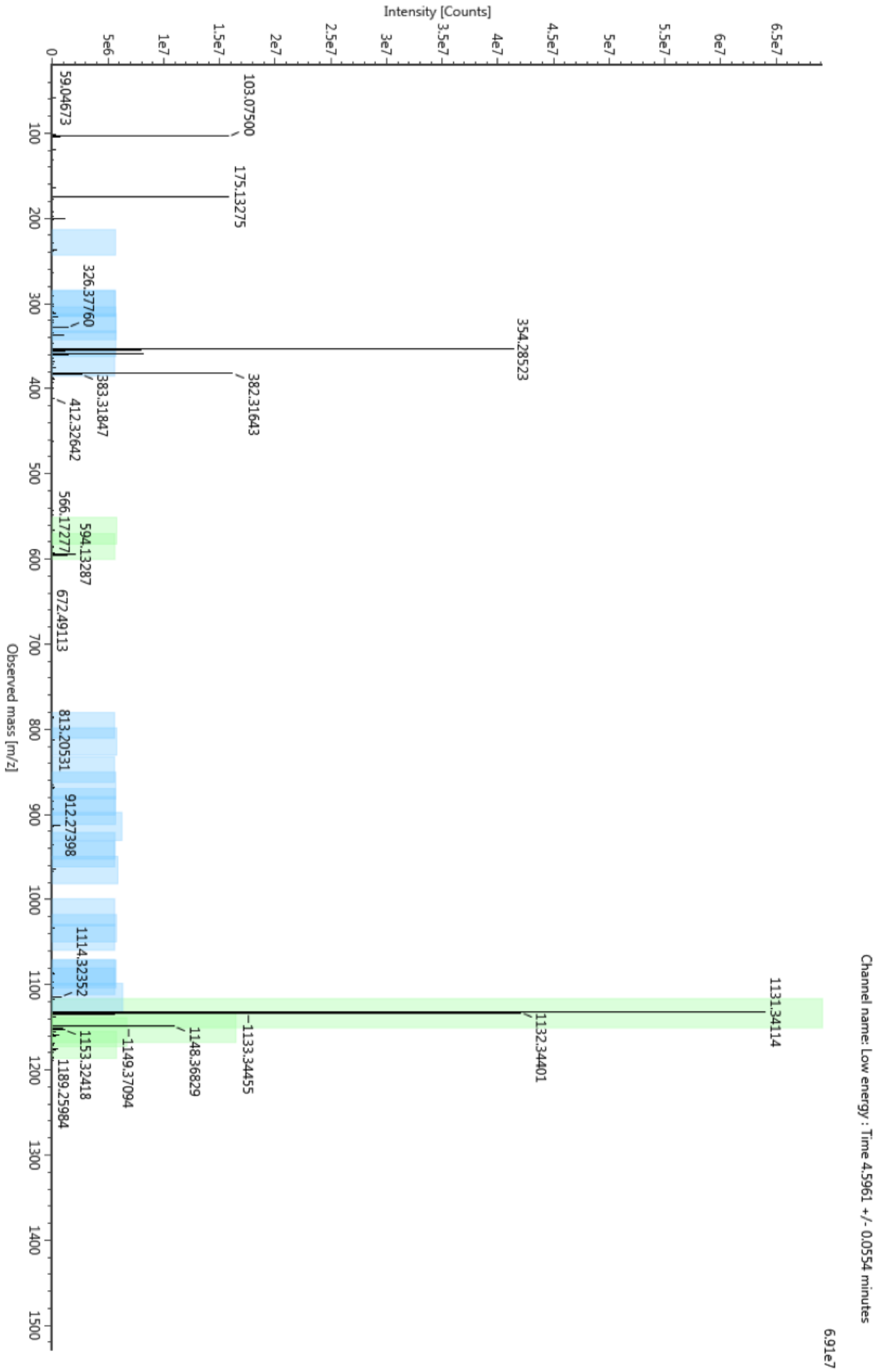


**Fig. S2.** Low energy mass spectrum for cutimycin ($[M+H]^+$ 1131.3411).

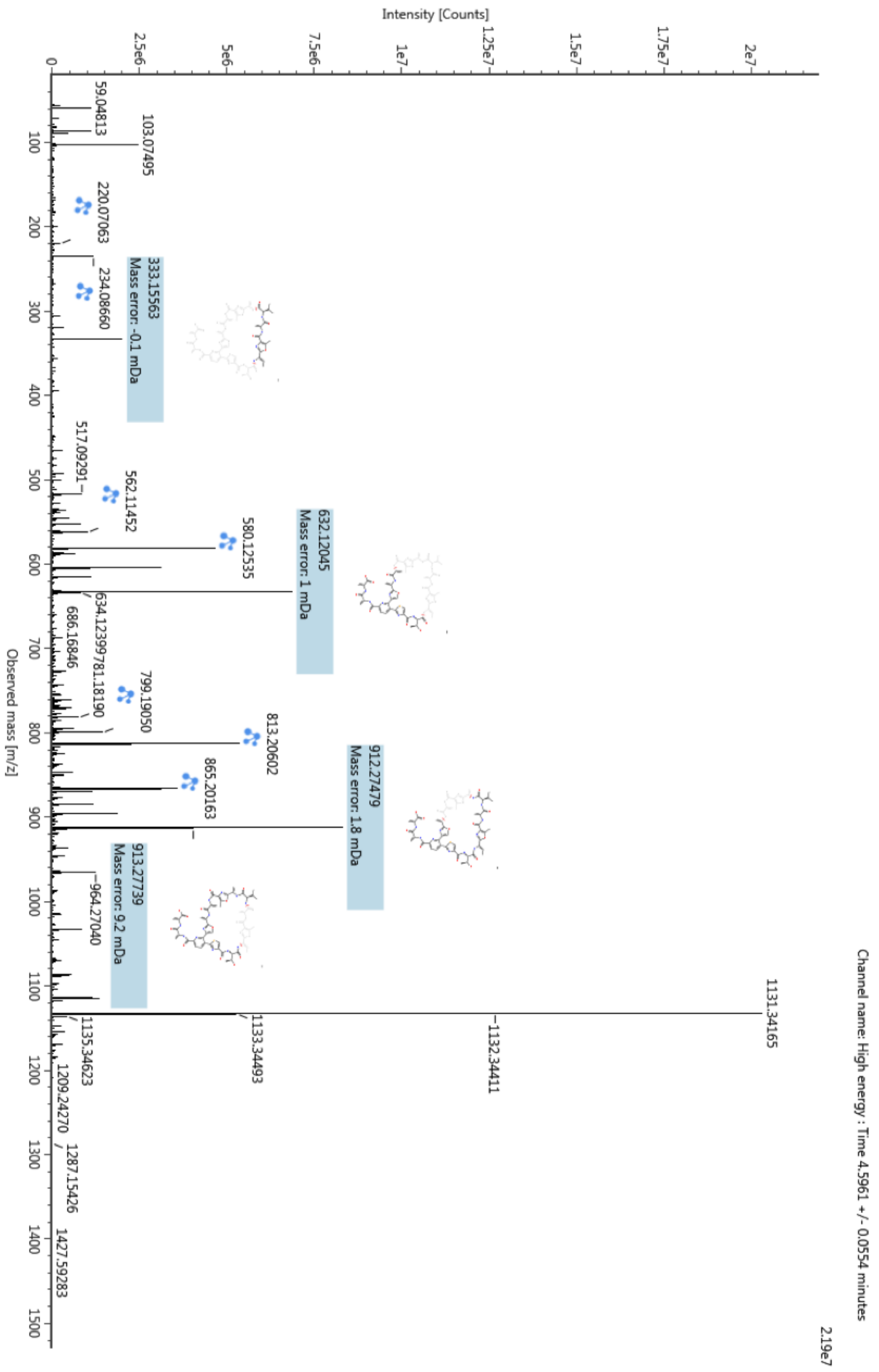


**Fig. S3.** High energy mass spectrum for cutimycin labeled with predicted fragments by Unifi.

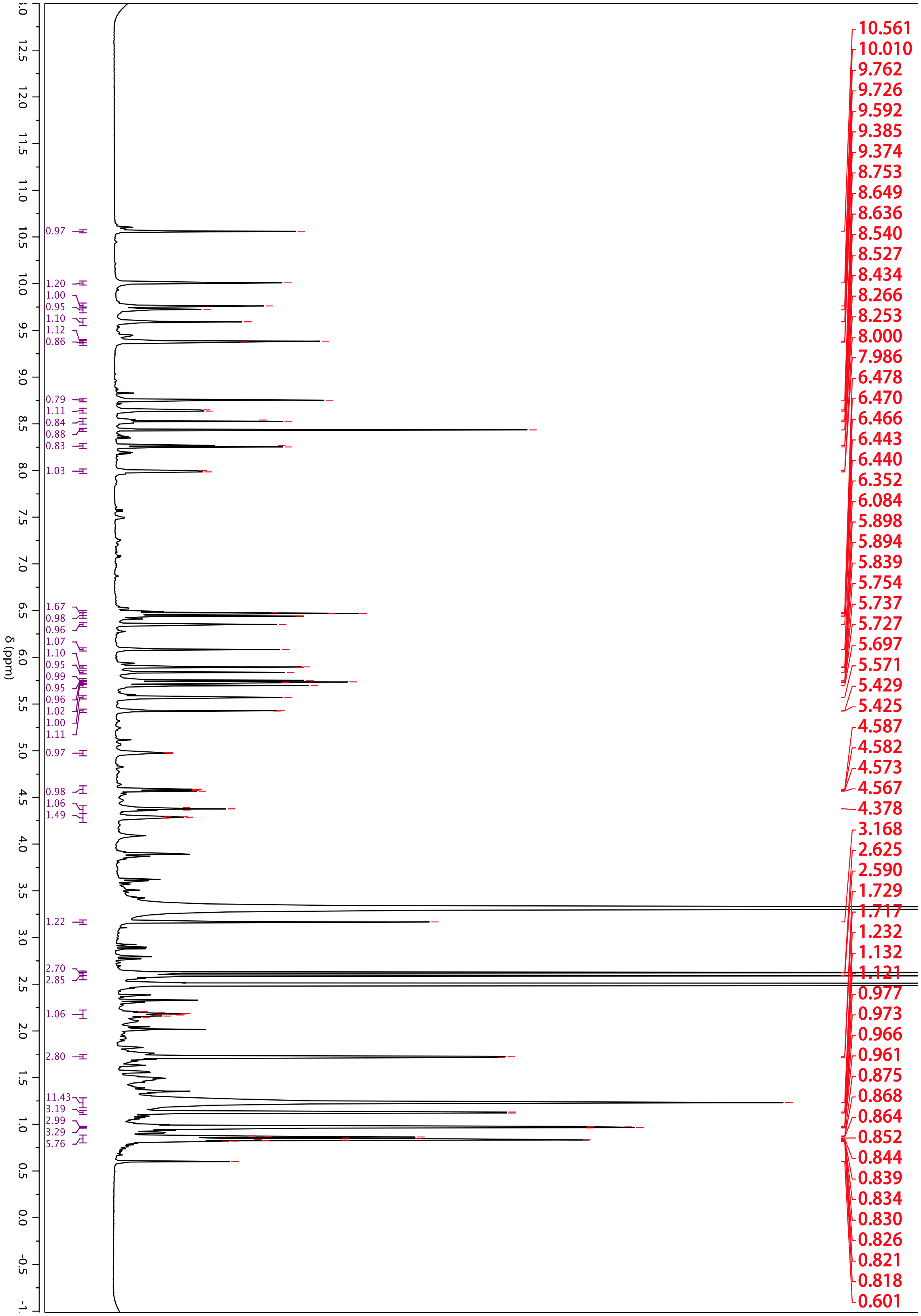

**Fig. S4.** [1]H NMR Spectrum of cutimycin taken in DMSO-*d6* at 600 MHz.

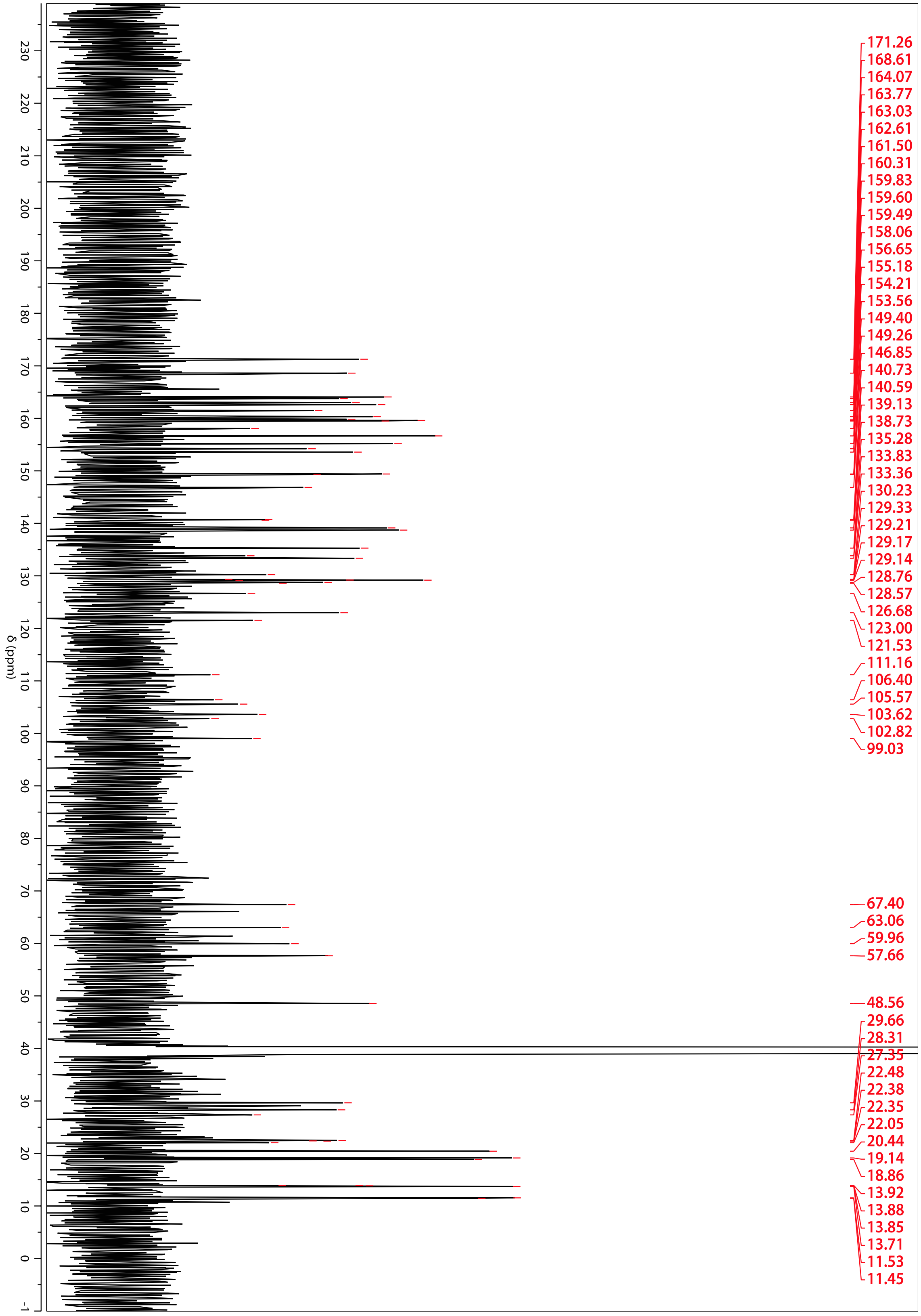


**Fig. S5.** $^{13}$C NMR Spectrum of cutimycin taken in DMSO-*d6* at 151 MHz.

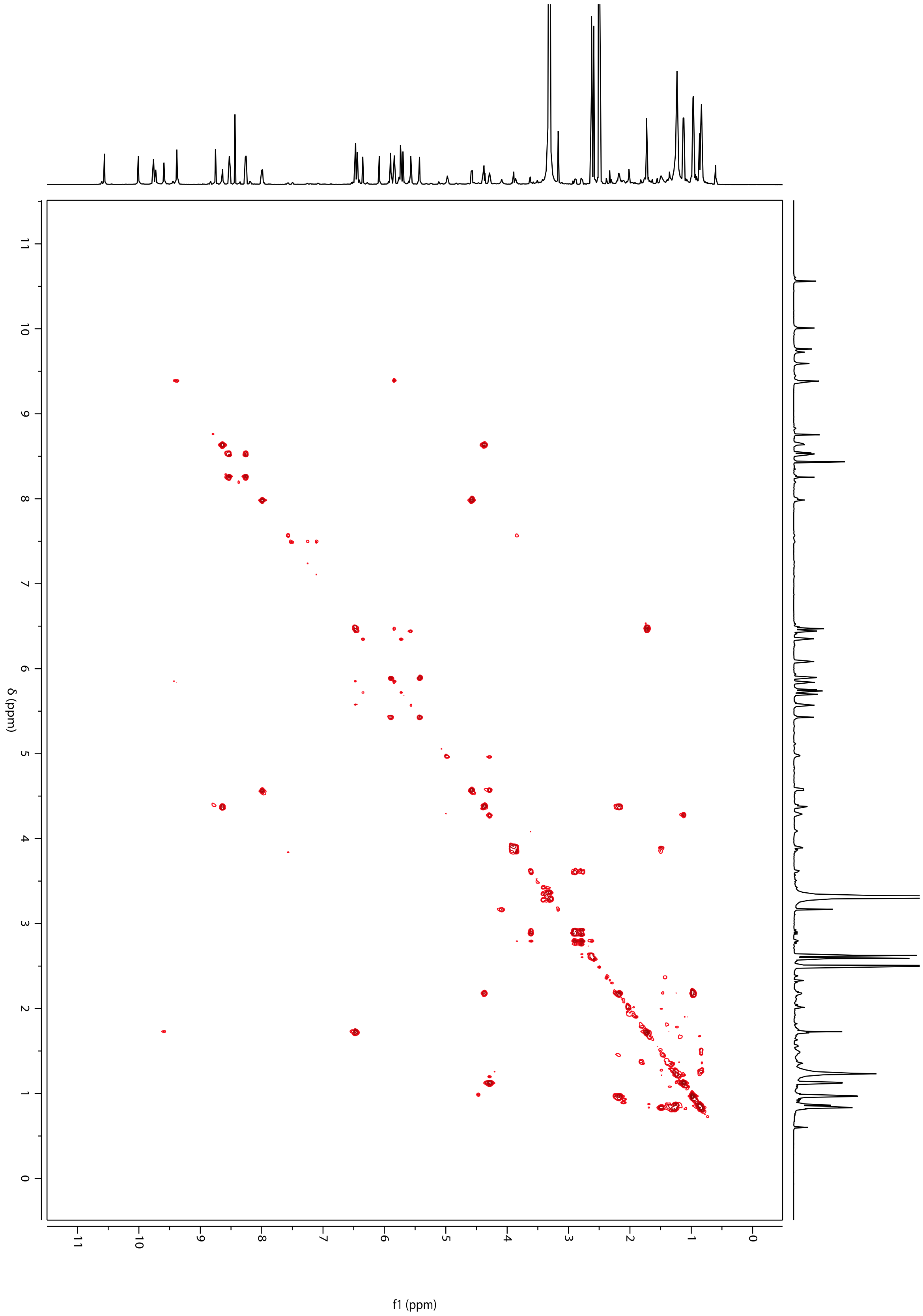

**Fig. S6.** ¹H-¹H COSY NMR in DMSO-*d6*.

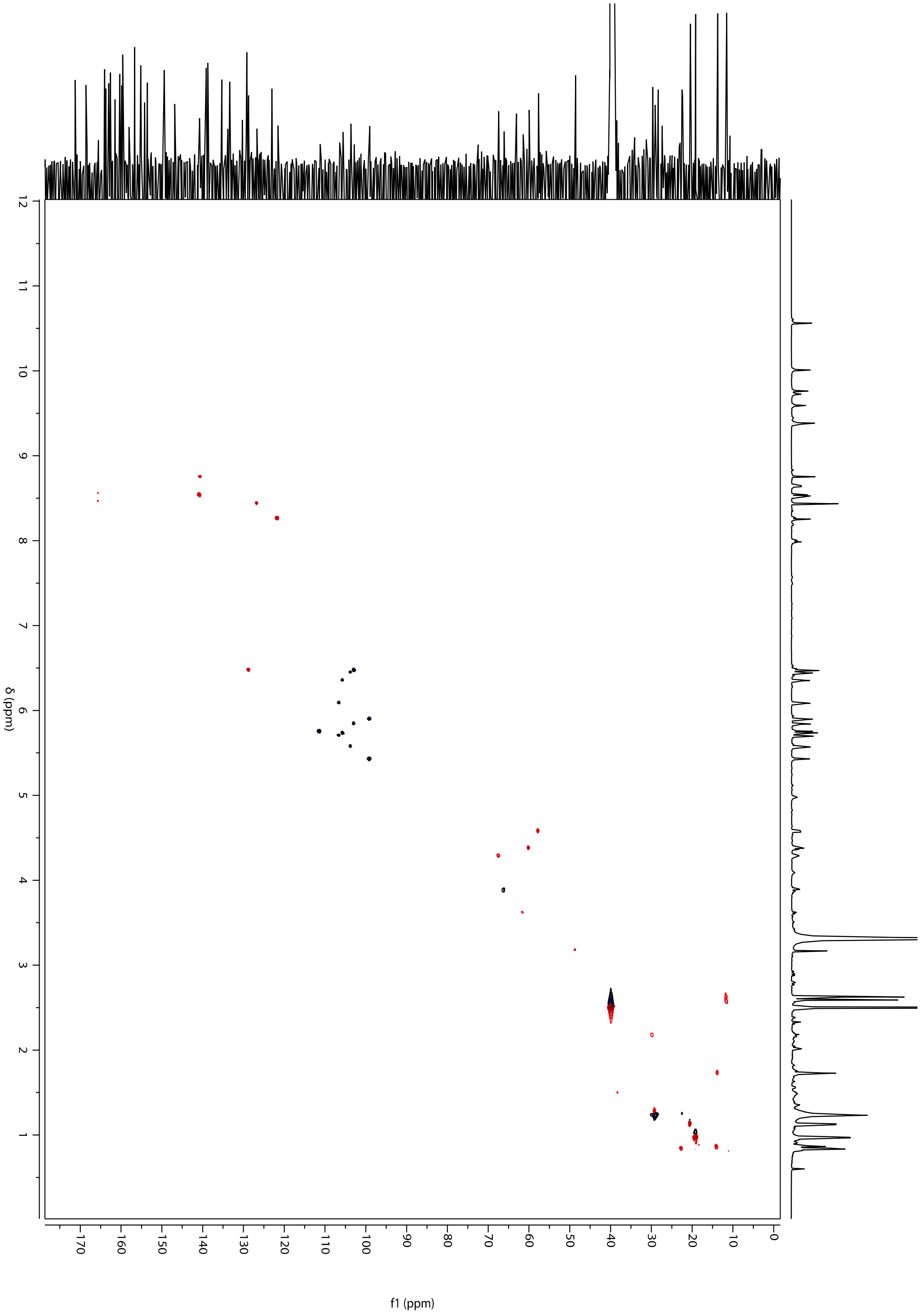


**Fig. S7.** $^{1}H$-$^{13}C$ HSQC NMR in DMSO-*d6*.

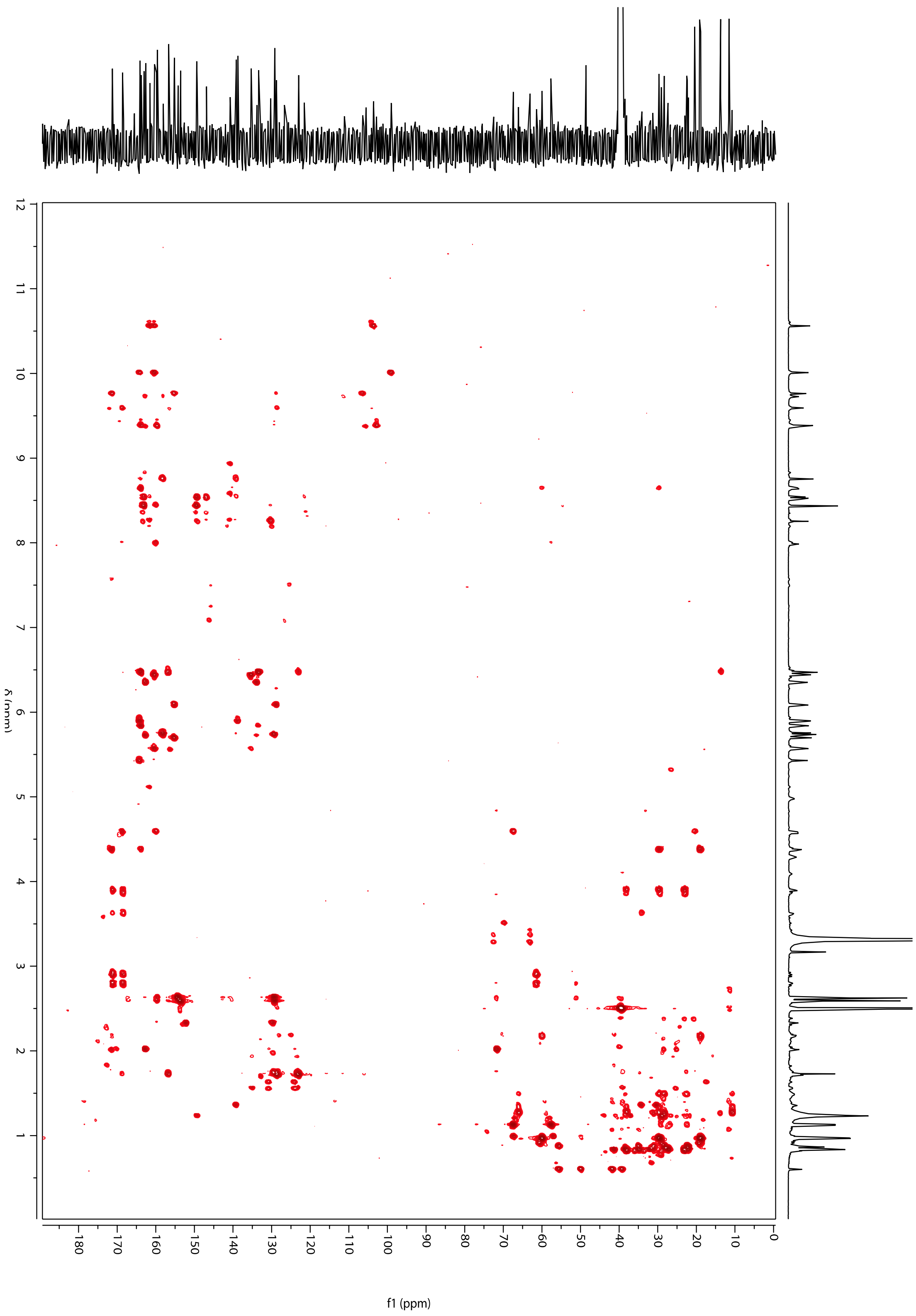

**Fig. S8.** $^{1}H$-$^{13}C$ HMBC NMR in DMSO-*d6*.

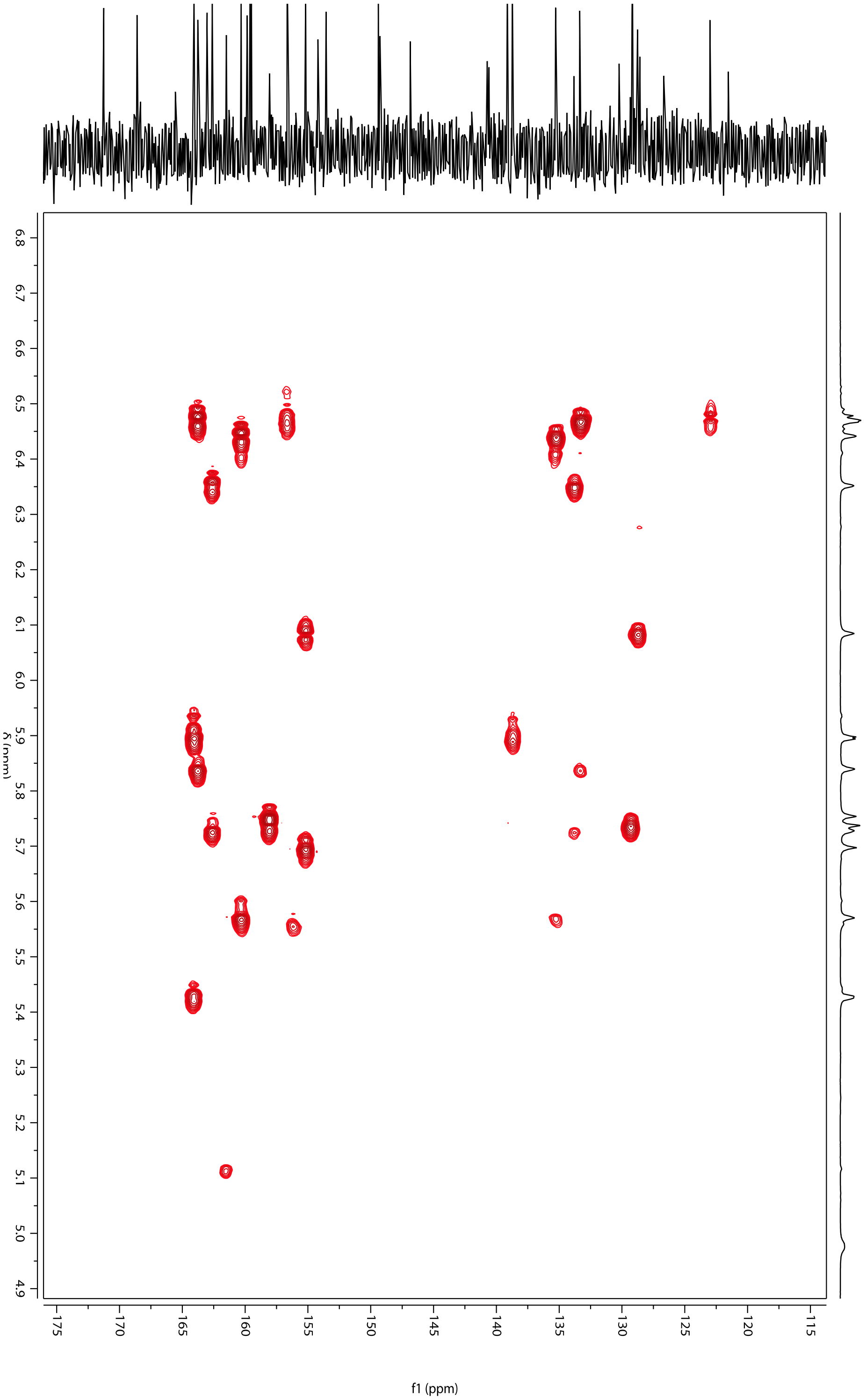


**Fig. S9.** Zoom in of $^{1}H$-$^{13}C$ HMBC NMR in DMSO-*d6* identifying dehydroalanine residues through presence of diastereotopic terminal alkene protons (position 20, $\delta_H$ 5.74, 5.75 and position 24 $\delta_H$ 5.73, 6.35) with HMBC correlations to carbon atoms 19 and 23 respectively.

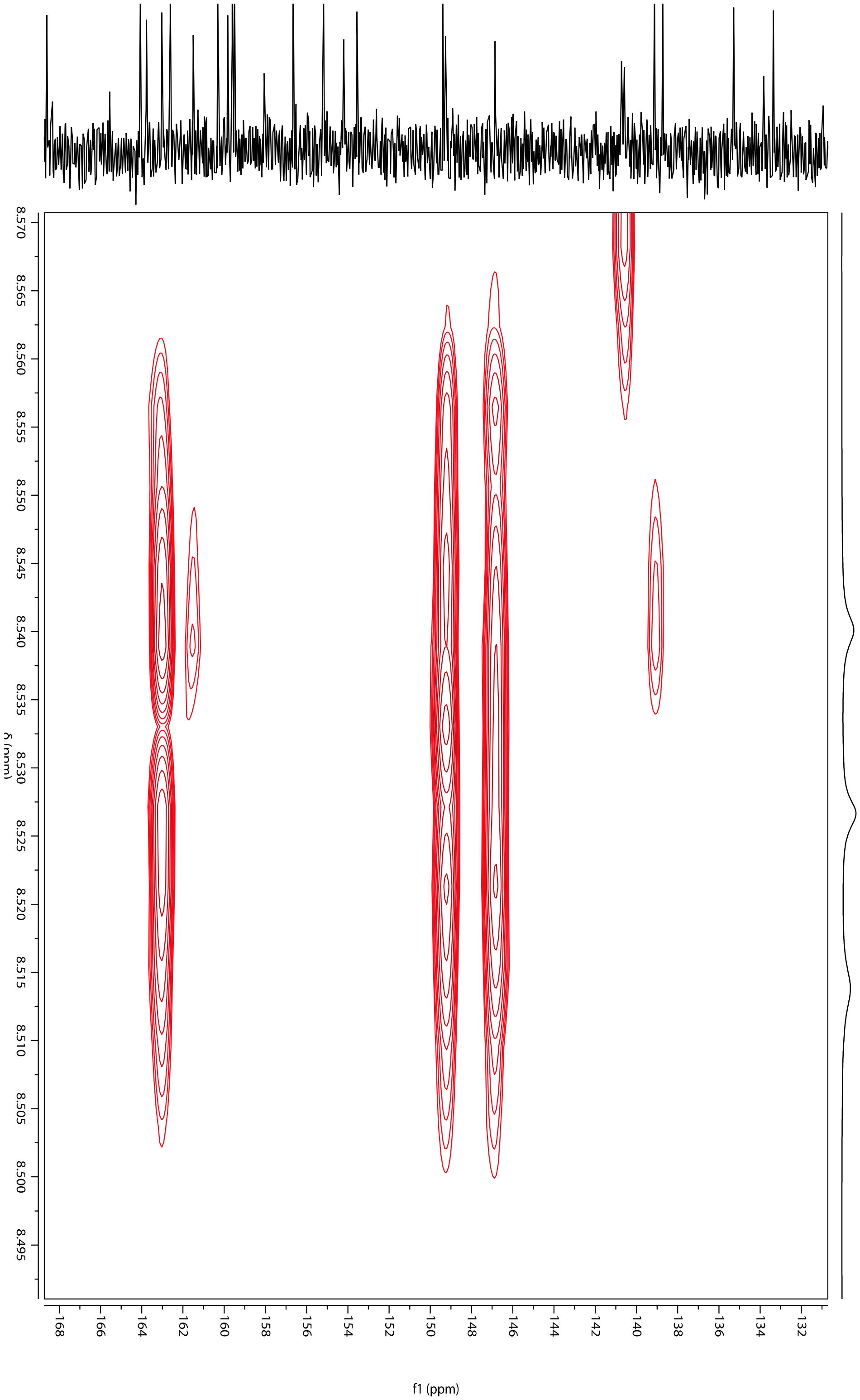


**Fig. S10.** Zoom in of $^{1}H$-$^{13}C$ HMBC NMR in DMSO-*d6* showing the weak four-bond HMBC correlation from position 13 to quaternary carbon 16 at $\delta_C$ 139. 13.

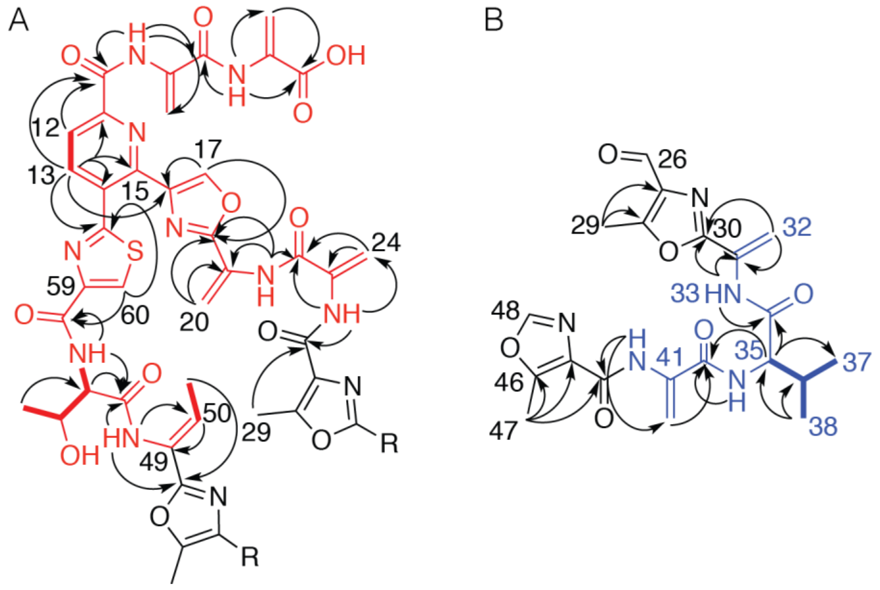


**FIG. S11.** The subunits of cutimycin with key HMBC (arrows) and COSY (bold bonds) correlations. A) Key HMBC correlations from the core of subunit A of cutimycin. B) Key HMBC correlations from subunit B of cutimycin.

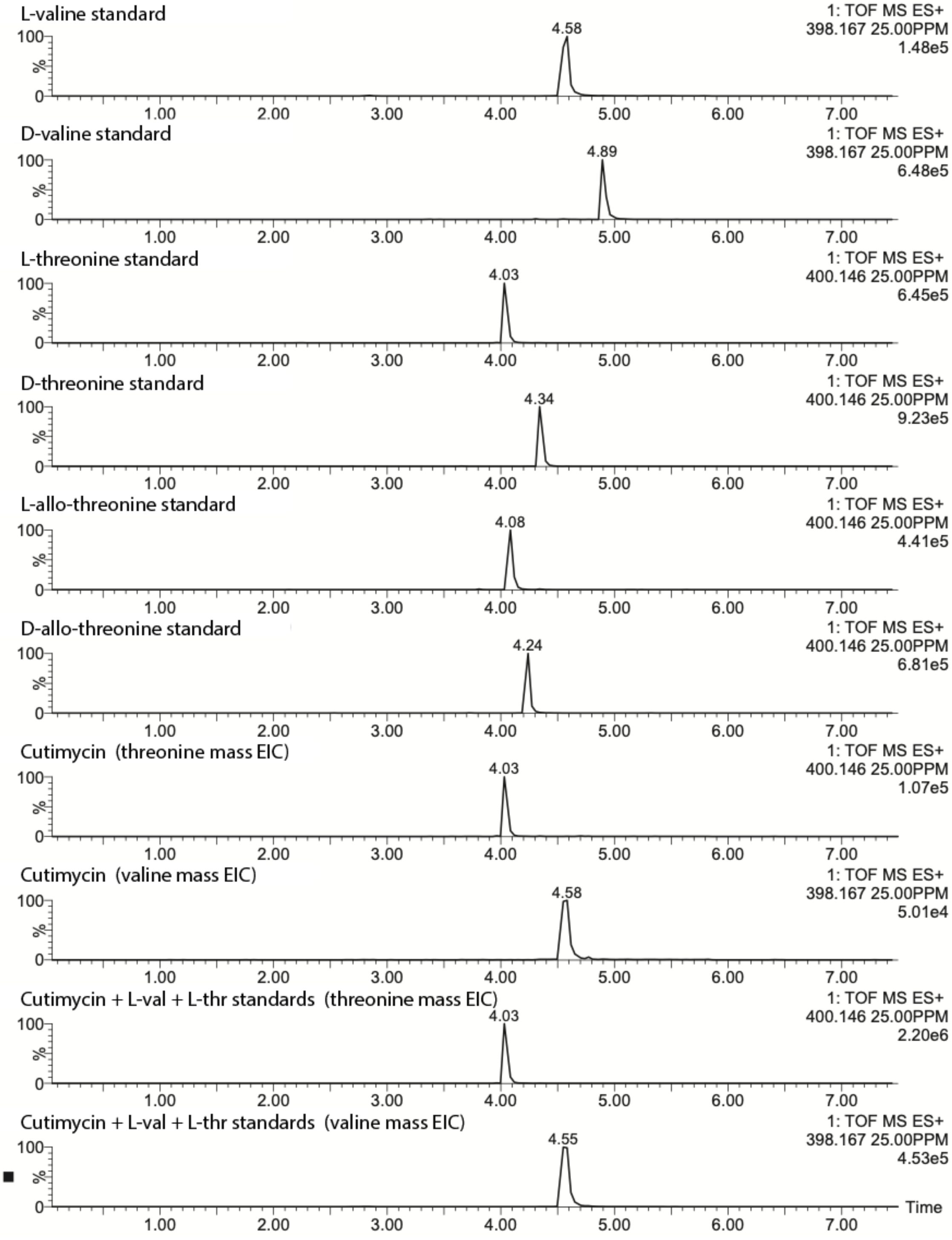


**Fig. S12.** Marfey's analysis of cutimycin. Individual amino acid Marfey's analysis derivatives, cutimycin hydrolysis product derivatives, and coinjections.

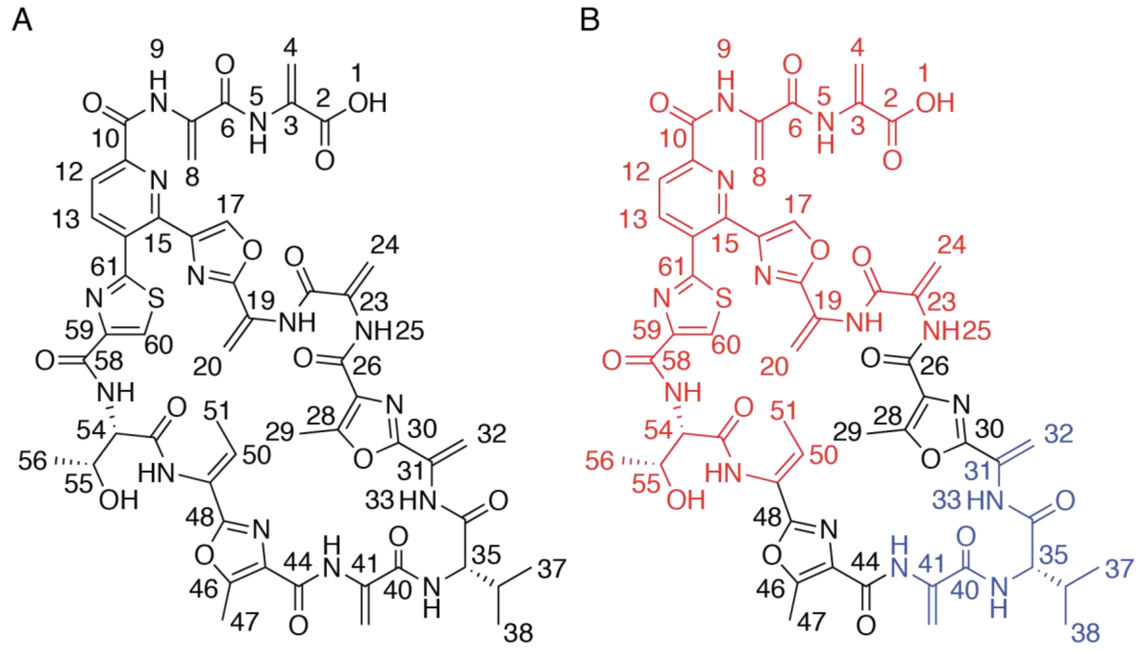


**Fig. S13.** Structure of cutimycin A) The full structure of cutimycin with atom positions labeled. B) The full structure of cutimycin showing subunits A (red) and B (blue) that were solved using NMR.

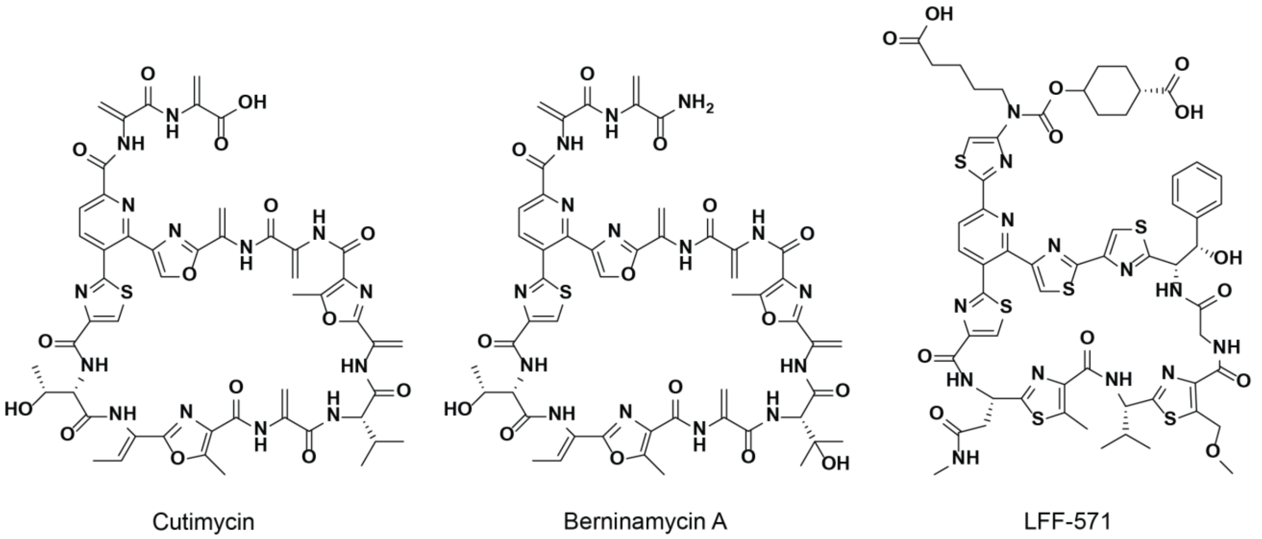


**Fig. S14.** The structures of A) cutimycin, B) berninamycin A, and C) LFF-571.

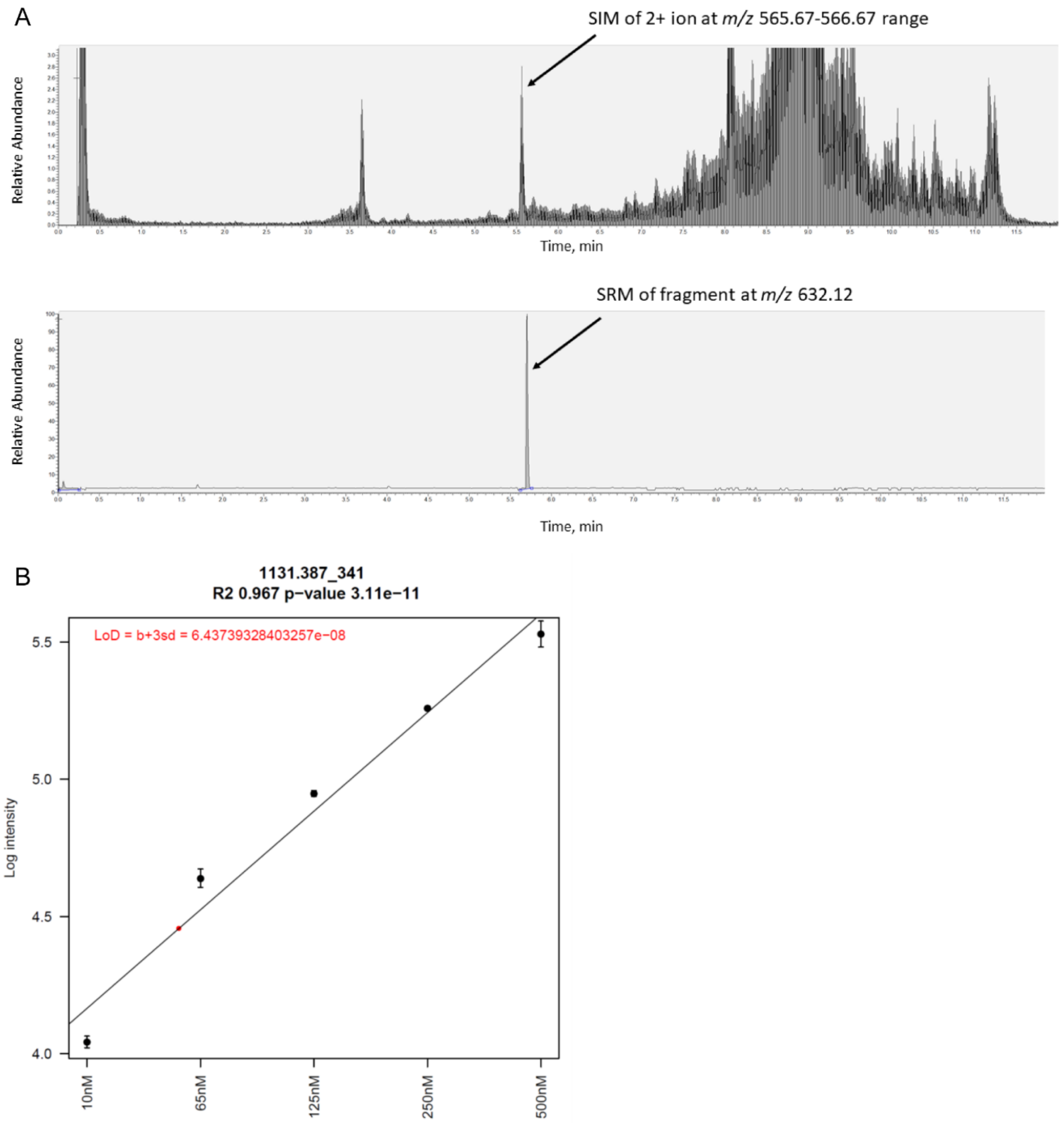


**Fig. S15.** Mass spectrometric detection of cutimycin from pooled human follicular content (n=60) for one of the tested samples. A) An example of mass spectrometric detection of cutimycin from pooled human follicular content for one of the tested samples. The top panel shows SIM of the 2+ ion of cutimycin; the bottom panel shows one of monitored MRM transitions of m/z 566 -> 632. Peak at the retention time corresponding to that of cutimycin standard are observed in both SIM and MRM. b) Calibration curve for the +1 ion of the cutimycin standard. The experimentally determined limit of detection in the extract is 64.4 nM.

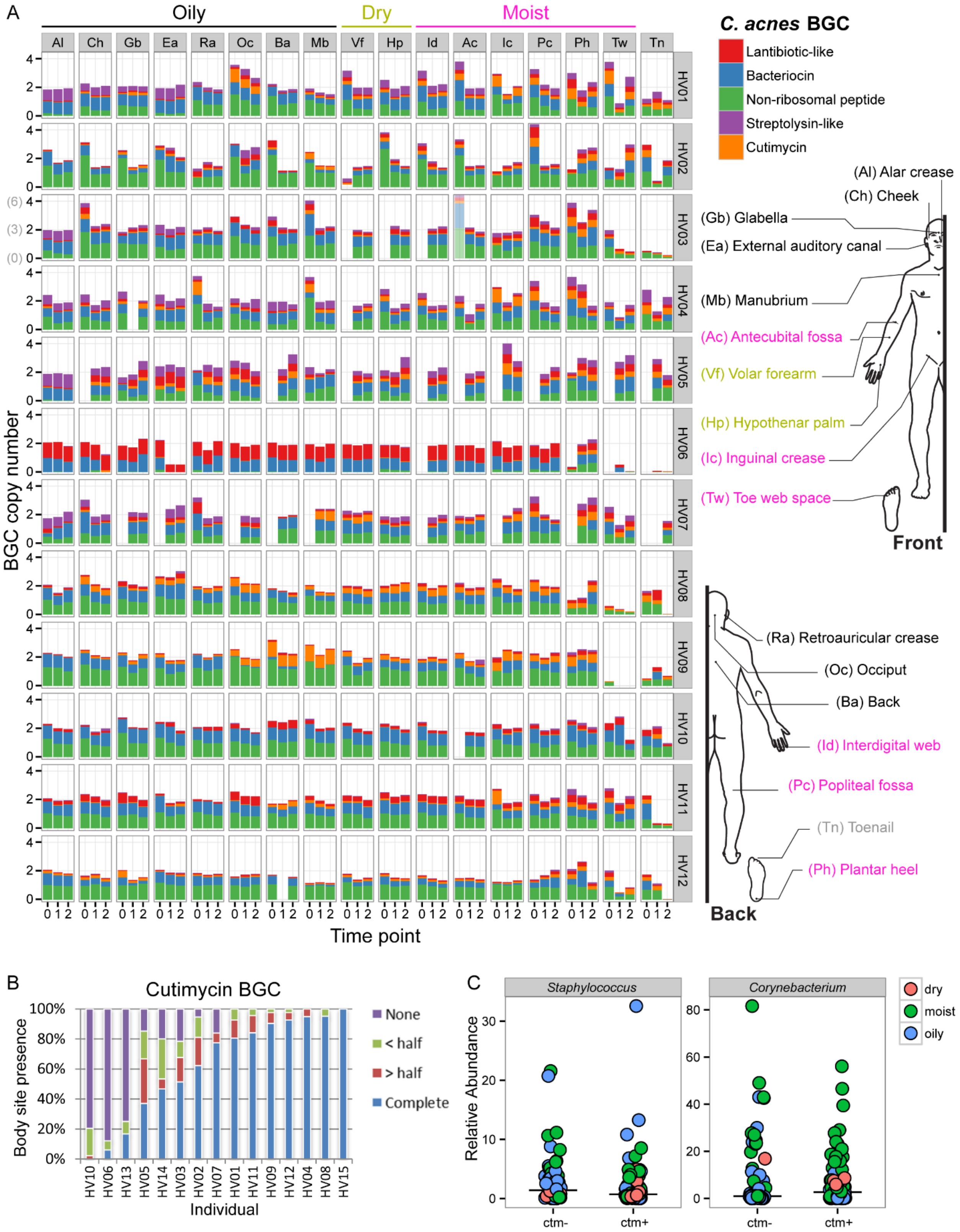


**Fig. S16.** Metagenomic analysis of *C. acnes* BGCs at 17 skin sites from sites from 12 healthy participants. A) Presence of *C. acnes* BGCs in the skin metagenomes from 12 healthy individuals over three time points. BGCs are color coded according to the legend (the cutimycin BGC is in orange) and relative copy numbers are stacked for each body site and time point. A BGC bar height of 1 indicates that particular BGC is present in ~100% of *C. acnes* at the particular sample site. Body sites are grouped according to their predominant oily, dry or moist characteristics, and abbreviations are explained in the inset. Note that for individual HV03, the Mean relative BGC abundance scale is depicted in grey, only for Ac at Time point 0. B) Percent of samples across body sites and time points from each participant for which the cutimycin BGC was detected in the metagenomic data. Separate genes are counted as present if over 40% of their length is covered by the sequence reads and the degree of genes detected for the cutimycin BGC is categorized as follows: "none" (purple), "less than half" (green), "greater than half" (red) and "complete" (blue). C) Metagenomics analysis did not reveal any correlation between the presence (ctm+) or absence (ctm-) of the cutimycin BGC with the genus *Corynebacterium* or the genus *Staphylococcus*. Ctm+ samples were those where all 9 genes of the cutimycin BGC were identified as present.

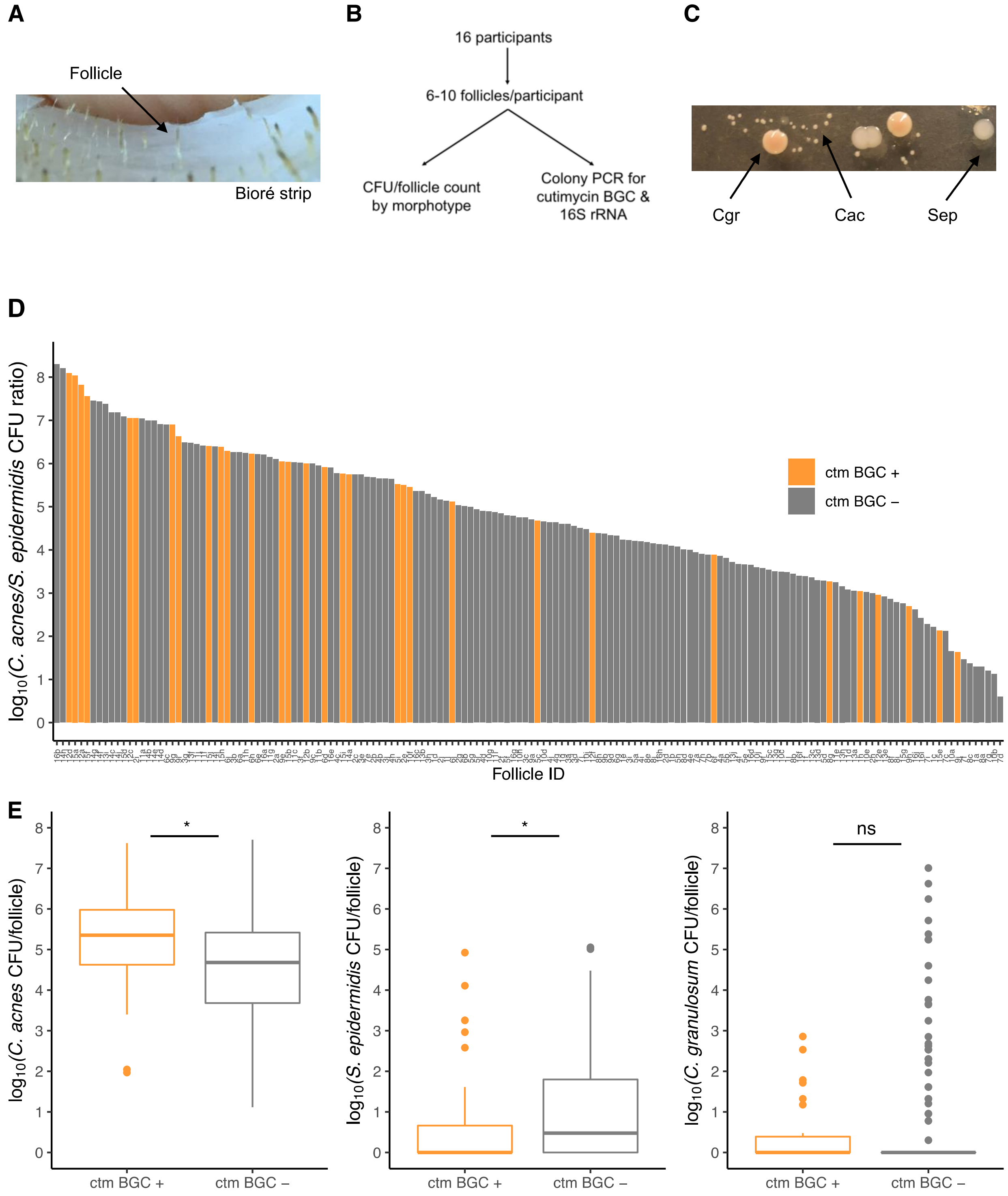


**Fig. S17.** The presence of the cutimycin BGC has a significant impact on the *C. acnes/S .epidermidis* ratio in human skin follicles. (**A**) Picture of Bioré pore strip with harvested follicular content. (**B**) Schematic of the experimental design. (**C**) Picture of culture isolates from follicular content showing the distinct colony morphologies of *C. acnes* (Cac), *S. epidermidis* (Sep) and *C. granulosum* (Cgr). **D**) Ranked histogram of the ratio of *C. acnes/S. epidermidis*

from each sampled follicle by the presence and absence of the cutimycin BGC. The Follicle ID denotes a participant by a number from 1-16 and pores by a letter from a-j (e.g. 10b). **E)** Box plots of the impact of the cutimycin BGC presence/absence on *C. acnes*, *C. granulosum* and *S. epidermidis* CFUs in human skin follicles. Data was pooled based on the assumption that follicles within an individual are independent and a non-parametric test (Wilcoxon signed-rank test) was used to compare the cutimycin BGC-positive and -negative PCR screen results. This showed that the CFUs of Cac ($p$=0.012) were significant, while the CFUs of Sep ($p$=0.042) and Cgr ($p$=0.979) were borderline or not significant, respectively, between cutimycin-positive and -negative samples.

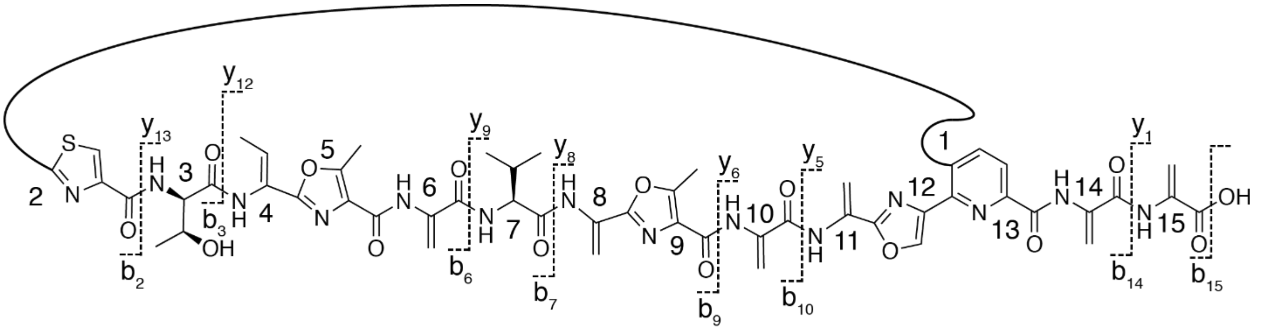

**Fig. S18.** Linear depiction of cutimycin with the key mass spec fragments labeled (Table S9).

| **Organism/Name** | **Strain** | **ppa0859 BLAST Results** |
|---|---|---|
| *Cutibacterium acnes* KPA171202 | KPA171202 | **Present** |
| *Cutibacterium acnes* 6609 | 6609 | **Present** |
| *Cutibacterium acnes* | PA_21_1_L1 | **Present** |
| *Cutibacterium acnes* | KCOM 1315 | **Present** |
| *Cutibacterium acnes* HL030PA1 | HL030PA1 | **Present** |
| *Cutibacterium acnes* | T14358 | **Present** |
| *Cutibacterium acnes* | Asn12 | **Present** |
| *Cutibacterium sp.* KPL2008 (putative *C. acnes*) | KPL2008 | **Present** |
| *Cutibacterium sp.* KPL2009 (putative *C. acnes*) | KPL2009 | Absent |
| *Cutibacterium sp.* KPL2005 (putative *C. avidum*) | KPL2005 | Absent |
| *Cutibacterium sp.* KPL1852 (putative *C. avidum*) | KPL1852 | Absent |
| *Cutibacterium sp.* KPL1844 (putative *C. granulosum*) | KPL1844 | Absent |
| *Cutibacterium sp.* KPL1838 (putative *C. avidum*) | KPL1838 | Absent |
| *Cutibacterium sp.* KPL2003 (putative *C. acnes*) | KPL2003 | Absent |
| *Cutibacterium sp.* KPL2000 (putative *C. avidum*) | KPL2000 | Absent |
| *Cutibacterium sp.* KPL1854 (putative *C. acnes*) | KPL1854 | Absent |
| *Cutibacterium sp.* KPL1849 (putative *C. acnes*) | KPL1849 | Absent |
| *Cutibacterium sp.* KPL1847 (putative *C. acnes*) | KPL1847 | Absent |
| *Cutibacterium avidum* 44067 | 44067 | Absent |
| *Cutibacterium avidum* | DPC 6544 | Absent |
| *Cutibacterium avidum* ATCC 25577 | ATCC 25577 | Absent |
| *Cutibacterium avidum* | MJR7694 | Absent |
| *Cutibacterium avidum* TM16 | TM16 | Absent |
| *Cutibacterium avidum* | UCD-PD2 | Absent |
| *Cutibacterium avidum* | T13 | Absent |
| *Cutibacterium avidum* | T15 | Absent |
| *Cutibacterium avidum* | T14 | Absent |
| *Cutibacterium avidum* | CI878 | Absent |
| *Cutibacterium avidum* | CI855 | Absent |
| *Cutibacterium avidum* | CI853 | Absent |
| *Cutibacterium avidum* | CI882 | Absent |
| *Cutibacterium avidum* | CI828 | Absent |
| *Cutibacterium avidum* | P16-029 | Absent |
| *Cutibacterium granulosum* DSM 20700 | DSM 20700 | Absent |
| *Cutibacterium granulosum* | NCTC11865 | Absent |
| *Cutibacterium granulosum* TM11 | TM11 | Absent |
| *Cutibacterium granulosum* DSM 20700 | DSM 20700 | Absent |
| *Cutibacterium acnes* SK137 | SK137 | Absent |

| | | |
|---|---|---|
| *Cutibacterium acnes* 266 | 266 | Absent |
| *Cutibacterium acnes subsp. defendens* ATCC 11828 | ATCC 11828 | Absent |
| *Cutibacterium acnes* TypeIA2 P.acn33 | P.acn33 | Absent |
| *Cutibacterium acnes* TypeIA2 P.acn17 | P.acn17 | Absent |
| *Cutibacterium acnes* TypeIA2 P.acn31 | P.acn31 | Absent |
| *Cutibacterium acnes* C1 | C1 | Absent |
| *Cutibacterium acnes* HL096PA1 | HL096PA1 | Absent |
| *Cutibacterium acnes* hdn-1 | hdn-1 | Absent |
| *Cutibacterium acnes* | KCOM 1861 (= ChDC B594) | Absent |
| *Cutibacterium acnes* | PA_15_1_R1 | Absent |
| *Cutibacterium acnes* | PA_15_2_L1 | Absent |
| *Cutibacterium acnes* | PA_12_1_L1 | Absent |
| *Cutibacterium acnes* | ATCC 6919 | Absent |
| *Cutibacterium acnes* | PA_30_2_L1 | Absent |
| *Cutibacterium acnes* | PA_12_1_R1 | Absent |
| *Cutibacterium acnes* | A1-14 | Absent |
| *Cutibacterium acnes* HL001PA1 | HL001PA1 | Absent |
| *Cutibacterium acnes* HL002PA1 | HL002PA1 | Absent |
| *Cutibacterium acnes* HL067PA1 | HL067PA1 | Absent |
| *Cutibacterium acnes* HL005PA4 | HL005PA4 | Absent |
| *Cutibacterium acnes* HL082PA2 | HL082PA2 | Absent |
| *Cutibacterium acnes* HL002PA2 | HL002PA2 | Absent |
| *Cutibacterium acnes* HL025PA2 | HL025PA2 | Absent |
| *Cutibacterium acnes* HL053PA2 | HL053PA2 | Absent |
| *Cutibacterium acnes* HL053PA1 | HL053PA1 | Absent |
| *Cutibacterium acnes* HL110PA1 | HL110PA1 | Absent |
| *Cutibacterium acnes* HL083PA1 | HL083PA1 | Absent |
| *Cutibacterium acnes* HL072PA1 | HL072PA1 | Absent |
| *Cutibacterium acnes* HL050PA1 | HL050PA1 | Absent |
| *Cutibacterium acnes* HL046PA1 | HL046PA1 | Absent |
| *Cutibacterium acnes* HL005PA1 | HL005PA1 | Absent |
| *Cutibacterium acnes* HL050PA3 | HL050PA3 | Absent |
| *Cutibacterium acnes* HL005PA3 | HL005PA3 | Absent |
| *Cutibacterium acnes* HL045PA1 | HL045PA1 | Absent |
| *Cutibacterium acnes* HL005PA2 | HL005PA2 | Absent |
| *Cutibacterium acnes* HL086PA1 | HL086PA1 | Absent |
| *Cutibacterium acnes* HL038PA1 | HL038PA1 | Absent |
| *Cutibacterium acnes* HL050PA2 | HL050PA2 | Absent |

| | | |
|---|---|---|
| *Cutibacterium acnes* HL087PA3 | HL087PA3 | Absent |
| *Cutibacterium acnes* HL020PA1 | HL020PA1 | Absent |
| *Cutibacterium acnes* HL013PA2 | HL013PA2 | Absent |
| *Cutibacterium acnes* HL063PA1 | HL063PA1 | Absent |
| *Cutibacterium acnes* HL036PA1 | HL036PA1 | Absent |
| *Cutibacterium acnes* HL036PA2 | HL036PA2 | Absent |
| *Cutibacterium acnes* HL027PA2 | HL027PA2 | Absent |
| *Cutibacterium acnes* HL063PA2 | HL063PA2 | Absent |
| *Cutibacterium acnes* HL043PA2 | HL043PA2 | Absent |
| *Cutibacterium acnes* HL002PA3 | HL002PA3 | Absent |
| *Cutibacterium acnes* HL025PA1 | HL025PA1 | Absent |
| *Cutibacterium acnes* HL110PA2 | HL110PA2 | Absent |
| *Cutibacterium acnes* HL074PA1 | HL074PA1 | Absent |
| *Cutibacterium acnes* HL059PA1 | HL059PA1 | Absent |
| *Cutibacterium acnes* HL046PA2 | HL046PA2 | Absent |
| *Cutibacterium acnes* HL110PA4 | HL110PA4 | Absent |
| *Cutibacterium acnes* HL110PA3 | HL110PA3 | Absent |
| *Cutibacterium acnes* HL056PA1 | HL056PA1 | Absent |
| *Cutibacterium acnes* HL087PA1 | HL087PA1 | Absent |
| *Cutibacterium acnes* HL083PA2 | HL083PA2 | Absent |
| *Cutibacterium acnes* HL007PA1 | HL007PA1 | Absent |
| *Cutibacterium acnes* HL043PA1 | HL043PA1 | Absent |
| *Cutibacterium acnes* HL072PA2 | HL072PA2 | Absent |
| *Cutibacterium acnes* HL060PA1 | HL060PA1 | Absent |
| *Cutibacterium acnes* HL082PA1 | HL082PA1 | Absent |
| *Cutibacterium acnes* HL092PA1 | HL092PA1 | Absent |
| *Cutibacterium acnes* HL027PA1 | HL027PA1 | Absent |
| *Cutibacterium acnes* HL059PA2 | HL059PA2 | Absent |
| *Cutibacterium acnes* HL036PA3 | HL036PA3 | Absent |
| *Cutibacterium acnes* HL030PA2 | HL030PA2 | Absent |
| *Cutibacterium acnes* HL078PA1 | HL078PA1 | Absent |
| *Cutibacterium acnes* HL037PA1 | HL037PA1 | Absent |
| *Cutibacterium acnes* HL013PA1 | HL013PA1 | Absent |
| *Cutibacterium acnes* HL087PA2 | HL087PA2 | Absent |
| *Cutibacterium acnes* HL096PA3 | HL096PA3 | Absent |
| *Cutibacterium acnes* HL096PA2 | HL096PA2 | Absent |
| *Cutibacterium acnes* HL103PA1 | HL103PA1 | Absent |
| *Cutibacterium acnes* HL097PA1 | HL097PA1 | Absent |
| *Cutibacterium acnes* HL099PA1 | HL099PA1 | Absent |

| | | |
|---|---|---|
| *Cutibacterium acnes* J139 | J139 | Absent |
| *Cutibacterium acnes* J165 | J165 | Absent |
| *Cutibacterium acnes* SK187 | SK187 | Absent |
| *Cutibacterium acnes* SK182 | SK182 | Absent |
| *Cutibacterium acnes* PRP-38 | PRP-38 | Absent |
| *Cutibacterium acnes* FZ1/2/0 | FZ1/2/0 | Absent |
| *Cutibacterium acnes* DSM 1897 | DSM 1897 | Absent |
| *Cutibacterium acnes* HL042PA3 | HL042PA3 | Absent |
| *Cutibacterium acnes* PA2 | PA2 | Absent |
| *Cutibacterium acnes* P6 | P6 | Absent |
| *Cutibacterium acnes* | HL411PA1 | Absent |
| *Cutibacterium acnes* | ATCC 6919 | Absent |
| *Cutibacterium acnes* HL202PA1 | HL202PA1 | Absent |
| *Cutibacterium acnes* | 50.1.L1 | Absent |
| *Cutibacterium acnes* | 51.1.L1 | Absent |
| *Cutibacterium acnes* | 49.1.L1 | Absent |
| *Cutibacterium acnes* | 32.1.L2 | Absent |
| *Cutibacterium acnes* | 46.1.L1 | Absent |
| *Cutibacterium acnes* | 37.1.L1 | Absent |
| *Cutibacterium acnes* | NTS_2003_1719 | Absent |
| *Cutibacterium acnes* | LRY_BL | Absent |
| *Cutibacterium acnes* | NTS_2004_10708 | Absent |
| *Cutibacterium acnes* | HB | Absent |
| *Cutibacterium acnes* | 09-23 | Absent |
| *Cutibacterium acnes subsp. defendens* | 09-323 | Absent |
| *Cutibacterium acnes subsp. acnes* | 12-89 | Absent |
| *Cutibacterium acnes subsp. defendens* | 10-482 | Absent |
| *Cutibacterium acnes* | 11-78 | Absent |
| *Cutibacterium acnes subsp. acnes* | 11-88 | Absent |
| *Cutibacterium acnes subsp. acnes* | 11-90 | Absent |
| *Cutibacterium acnes subsp. defendens* | 10-43 | Absent |
| *Cutibacterium acnes* | 09-263 | Absent |
| *Cutibacterium acnes subsp. defendens* | 09-109 | Absent |
| *Cutibacterium acnes subsp. acnes* | 10-113 | Absent |
| *Cutibacterium acnes subsp. acnes* | 10-118 | Absent |
| *Cutibacterium acnes* | 10-167 | Absent |
| *Cutibacterium acnes subsp. defendens* | 11-49 | Absent |
| *Cutibacterium acnes subsp. defendens* | 11-79 | Absent |
| *Cutibacterium acnes subsp. defendens* | 11-356 | Absent |

| | | |
|---|---|---|
| *Cutibacterium acnes* | 09-9 | Absent |
| *Cutibacterium acnes* | CA17 | Absent |
| *Cutibacterium acnes* | CA39 | Absent |
| *Cutibacterium acnes* | CA51 | Absent |
| *Cutibacterium acnes* | 09-193 | Absent |
| *Cutibacterium acnes* | 09-322 | Absent |
| *Cutibacterium acnes* | 09-29 | Absent |
| *Cutibacterium acnes* | 09-258 | Absent |
| *Cutibacterium acnes* | UMB0211 | Absent |
| *Cutibacterium acnes* | P15-231 | Absent |
| *Cutibacterium acnes* | P15-207 | Absent |
| *Cutibacterium acnes* | P15-206 | Absent |
| *Cutibacterium acnes* | P15-180 | Absent |
| *Cutibacterium acnes* | P15-178 | Absent |
| *Cutibacterium acnes* | M13605 | Absent |
| *Cutibacterium acnes* | P15-186 | Absent |
| *Cutibacterium acnes* | T14441 | Absent |
| *Cutibacterium acnes* | T14475 | Absent |
| *Cutibacterium acnes* | T16975 | Absent |
| *Cutibacterium acnes* | T17001 | Absent |
| *Cutibacterium acnes* | T17039 | Absent |
| *Cutibacterium acnes* | T20574 | Absent |
| *Cutibacterium acnes* | T20670 | Absent |
| *Cutibacterium acnes* | T20736 | Absent |
| *Cutibacterium acnes* | T20758 | Absent |
| *Cutibacterium acnes* | T20816 | Absent |
| *Cutibacterium acnes* | T29350 | Absent |
| *Cutibacterium acnes* | T29362 | Absent |
| *Cutibacterium acnes* | T29420 | Absent |
| *Cutibacterium acnes* | T32516 | Absent |
| *Cutibacterium acnes* | T35709 | Absent |
| *Cutibacterium acnes* | T35743 | Absent |
| *Cutibacterium acnes* | T36318 | Absent |
| *Cutibacterium acnes* | P16-206 | Absent |
| *Cutibacterium acnes* | P15-077 | Absent |
| *Cutibacterium acnes* | P15-071 | Absent |
| *Cutibacterium acnes* | P15-014 | Absent |
| *Cutibacterium acnes* | P15-021 | Absent |
| *Cutibacterium acnes* | P15-089 | Absent |

| | | |
|---|---|---|
| *Cutibacterium acnes* | P15-159 | Absent |
| *Cutibacterium acnes* | P15-165 | Absent |
| *Cutibacterium acnes* | T14076 | Absent |
| *Cutibacterium acnes* | NLAE-zl-G260 | Absent |
| *Cutibacterium acnes* | 523_PAVI | Absent |
| *Cutibacterium acnes* | 52.1.L4 | Absent |
| *Cutibacterium acnes* | 48.1.L1 | Absent |
| *Cutibacterium acnes* | 26.1.L1 | Absent |
| *Cutibacterium acnes* | 43.1.L1 | Absent |
| *Cutibacterium acnes* | 44.1.L1 | Absent |
| *Cutibacterium acnes* | NTS_31306190 | Absent |
| *Cutibacterium acnes* JCM 18909 | JCM 18909 | Absent |
| *Cutibacterium acnes* JCM 18916 | JCM 18916 | Absent |
| *Cutibacterium acnes* JCM 18918 | JCM 18918 | Absent |
| *Cutibacterium acnes* JCM 18920 | JCM 18920 | Absent |
| *Cutibacterium acnes* HL201PA1 | HL201PA1 | Absent |
| *Cutibacterium acnes* | JCM 18919 | Absent |
| *Cutibacterium acnes* | S2_005_002_R2_31 | Absent |
| *Cutibacterium acnes* | T35877 | Absent |
| *Cutibacterium acnes* | 119_PAVI | Absent |
| *Cutibacterium acnes* | PMH5 | Absent |
| *Cutibacterium acnes* | PMH7 | Absent |
| *Cutibacterium acnes* | UBA1564 | Absent |
| *Cutibacterium acnes* | UBA3960 | Absent |
| *Cutibacterium acnes* | UBA11121 | Absent |
| *Cutibacterium acnes* | UBA9075 | Absent |

**Table S1.** Cutimycin BGC presence/absence in 219 *Cutibacterium* genomes

| Atom# | $^1$H | $^{13}$C | Integral | Multiplet | COSY | HMBC |
|---|---|---|---|---|---|---|
| 1 | - | OH | - | - | - | - |
| 2 | - | 164.07 | - | - | - | - |
| 3 | - | 138.73 | - | - | - | - |
| 4 | 5.43 | 99.03 | 1.00 | d, *J* = 2.5 Hz | 10.01, 5.90 | 164.07 |
| 4 | 5.9 | 99.03 | 1.00 | d, *J* = 2.4 Hz | 5.43 | 164.07, 138.73 |
| 5 | 10.01 | NH | 1.00 | S | 5.43 | 164.07, 160.31, 99.03 |
| 6 | - | 160.31 | - | - | - | - |
| 7 | - | 135.28 | - | - | - | - |
| 8 | 5.57 | 103.62 | 1.00 | S | 10.56, 6.44 | 160.31 |
| 8 | 6.44 | 103.62 | 1.00 | S | - | 160.31, 135.28 |
| 9 | 10.56 | NH | 1.00 | S | 5.57 | 161.50, 160.31, 103.62 |
| 10 | - | 161.5 | - | - | - | - |
| 11 | - | 149.26[a] | - | - | - | - |
| 12 | 8.26 | 121.53 | 1.00 | d, *J* = 8.1 Hz | 8.53 | 161.50, 140.73, 130.23 |
| 13 | 8.53 | 140.73 | 1.00 | d, *J* = 8.1 Hz | 8.26 | 163.03, 161.50, 149.26, 146.85, 139.13, 121.53 |
| 14 | - | 130.23[a] | - | - | - | - |
| 15 | - | 146.85[a] | - | - | - | - |
| 16 | - | 139.13 | - | - | - | - |
| 17 | 8.75 | 140.59 | 1.00 | S | - | 158.06, 139.13 |
| 18 | - | 158.06 | - | - | - | - |
| 19 | - | 129.33 | - | - | - | - |
| 20 | 5.74 | 111.16 | 1.00 | S | - | 158.06, 129.33 |
| 20 | 5.75 | 111.16 | 1.00 | S | - | 158.06, 129.33 |
| 21 | 9.73 | NH | 1.00 | S | - | 162.61, 158.06, 111.16 |
| 22 | - | 162.61 | - | - | - | - |
| 23 | - | 133.83 | - | - | - | - |
| 24 | 5.73 | 105.57 | 1.00 | S | 6.35 | 162.61, 133.83 |
| 24 | 6.35 | 105.57 | 1.00 | S | 5.73 | 162.61, 133.83 |
| 25 | 9.37 | NH | 1.00 | S | 5.73 | 162.61, 159.49,105.57 |
| 26 | - | 159.49 | - | - | - | - |
| 27 | - | 154.21[b] | - | - | - | - |
| 28 | - | 129.17[b] | - | - | - | - |
| 29 | 2.62 | 11.45 | 3.00 | S | - | 159.49, 154.21, 129.17 |
| 30 | - | 155.18 | - | - | - | - |
| 31 | - | 128.76 | - | - | - | - |
| 32 | 5.7 | 106.4 | 1.00 | S | 9.76 | 155.18, 128.76 |

| | | | | | | |
|---|---|---|---|---|---|---|
| 32 | 6.08 | 106.4 | 1.00 | S | - | 155.18, 128.76 |
| 33 | 9.76 | NH | 1.00 | S | 5.7 | 171.26, 155.18, 106.4 |
| 34 | - | 171.26 | - | - | - | - |
| 35 | 4.38 | 59.96 | 1.00 | t, *J* = 8.2, 8.2 Hz | 8.64, 2.18 | 171.26, 163.77 |
| 36 | 2.18 | 29.66 | 1.00 | dtd, *J* = 4.4, 7.8, 8.7, 15.5 Hz | 4.38, 0.97 | 171.26, 18.86 |
| 37,38 | 0.96 | 18.86 | 3.00 | d, *J* = 2.4 Hz | 2.18 | 59.96, 29.66 |
| 37,38 | 0.97 | 19.14 | 3.00 | d, *J* = 2.4 Hz | 2.18 | 59.96, 29.66 |
| 39 | 8.64 | NH | 1.00 | d, *J* = 7.7 Hz | 4.38 | 163.77, 59.96, 29.7 |
| 40 | - | 163.77 | - | - | - | - |
| 41 | - | 133.36 | - | - | - | - |
| 42 | 5.84 | 102.82 | 1.00 | S | - | 163.77, 133.36 |
| 42 | 6.46 | 102.82 | 1.00 | d, *J* = 7.4 Hz | 5.84, 1.72 | 163.77, 156.65, 133.36 |
| 43 | 9.39 | NH | 1.00 | S | 5.84 | 163.77, 159.6, 102.82 |
| 44 | - | 159.60 | - | - | - | - |
| 45 | - | 153.56[c] | - | - | - | - |
| 46 | - | 129.14[c] | - | - | - | - |
| 47 | 2.59 | 11.53 | 3.00 | S | - | 159.60, 153.56, 129.14 |
| 48 | - | 156.65 | - | - | - | - |
| 49 | - | 123.00 | - | - | - | - |
| 50 | 6.47 | 128.57 | 1.00 | d, *J* = 5.0 Hz | 1.72 | 123.0, 13.71 |
| 51 | 1.72 | 13.71 | 3.00 | d, *J* = 7.2 Hz | 6.47 | 156.65, 128.57, 123.0 |
| 52 | 9.59 | NH | 1.00 | s | - | 168.61, 156.65, 128.57 |
| 53 | - | 168.61 | - | - | - | - |
| 54 | 4.58 | 57.66 | 1.00 | dd, *J* = 3.3, 8.8 Hz | 7.99, 4.29 | 168.61, 159.83, 67.4, 20.44 |
| 55 | 4.29 | 67.40 | 1.00 | d, *J* = 6.5 Hz | 4.58, 4.98, 1.13 | - |
| 56 | 1.13 | 20.44 | 3.00 | d, *J* = 6.3 Hz | 4.29 | 57.66, 67.4 |
| 57 | 7.99 | NH | 1.00 | d, *J* = 8.7 Hz | 4.58 | 168.61, 159.83 |
| 58 | - | 159.83 | - | - | - | - |
| 59 | - | 149.40 | - | - | - | - |
| 60 | 8.43 | 126.68 | 1.00 | s | - | 163.03, 159.83, 149.40 |
| 61 | - | 163.03 | - | - | - | - |
| 62 | 4.98 | OH | 1.00 | d, *J* = 5.6 Hz | 4.29 | - |

**Table S2:** NMR Data of cutimycin taken in DMSO-*d6* at 600 MHz and 151 MHz.
[a, b, c]Assignments interchangable

| Strain | cutimycin | berninamycin |
|---|---|---|
| *C. acnes* KPA171202* | >3.2 | >3.2 |
| *C. acnes* HL030PA1* | >3.2 | >3.2 |
| *C. acnes* HL110PA1 | >3.2 | >3.2 |
| *C. acnes* HL086PA1 | >3.2 | >3.2 |
| *C. acnes* HL053PA2 | >3.2 | >3.2 |
| *S. aureus* USA300 NRS384 | 0.2 | 0.8 |
| *S. aureus* UAMS-1 | 0.8 | 3.2 |
| *S. epidermidis* W23144 | 0.2 | 0.8 |
| *S. epidermidis* DSM20042 | 0.8 | 3.2 |
| *S. epidermidis* ATCC 35984 | 3.2 | 3.2 |
| *C. accolens* ATCC 49725 | >3.2 | >3.2 |
| *C. jeikeium* DSM7171 | >3.2 | >3.2 |
| *C. striatum* DSM20668 | 3.2 | 3.2 |
| *C. pseudodiphtheriticum* DSM44287 | >3.2 | >3.2 |

**Table S3.** Minimal inhibitory concentrations for cutimycin and bernamycin measured at 16 hours of growth for skin- and nasal-associated *Staphylococcus* and *Corynebacterium* species and 3 days for *BGC+ and BGC- *C. acnes*. Concentrations tested (μM): 0, 0.05, 0.2, 0.8, 3.2.

| **Sample ID, experiment 1** | **Cutimycin detected** | **Sample ID, experiment 2** | **Cutimycin detected** |
|---|---|---|---|
| Tube 1 | No | Subject 1 (pool, 35 follicles) | No |
| Tube 2 | Yes | Subject 2 (pool, 80 follicles) | No |
| Tube 3 | Yes | Subject 3 (pool, 55 follicles) | No |
| Tube 4 | No | Subject 4 (pool, 60 follicles) | Yes |
| Tube 5 | No | Subject 5 (pool, 25 follicles) | No |
| Tube 6 | No | | |
| Tube 7 | No | | |
| Tube 8 | Yes | | |
| Tube 9 | Yes | | |
| Tube 10 | No | | |
| Tube 11 | No | | |
| Tube 12 | No | | |
| Tube 13 | No | | |
| Tube 14 | Yes | | |
| Tube 15 | Yes | | |
| Tube 16 | No | | |
| Tube 17 | No | | |
| Tube 18 | No | | |
| Tube 19 | No | | |
| Tube 20 | No | | |
| Tube 21 | No | | |
| Tube 22 | Yes | | |
| Tube 23 | Yes | | |
| Tube 24 | No | | |
| Tube 25 | No | | |
| Tube 26 | No | | |
| Tube 27 | No | | |
| Tube 28 | Yes | | |

**Table S4.** Cutimycin in human follicular samples has been detected in 28% of all samples across two separate experiments.

| BGC type (representative strain) | Locus tags |
| --- | --- |
| Bacteriocin - Lactococcin 972-like (KPA171202) | PPA_RS04095 |
| Non-ribosomal peptide (KPA171202) | PPA_RS12620 – PPA_RS06510 |
| Cutimycin (KPA171202) | PPA_RS13410 – PPA_RS12420 |
| Lantibiotic-like (HL053PA2) | HMPREF9565_00657 – _00677 |
| Streptolysin-like (SK137) | HMPREF0675_3180 – _3189 |

**Table S5.** Locus tags for the *C. acnes* BGCs assayed for in skin metagenomic data.

| Follicle ID | *C. acnes* CFUs | *C. granulosum* CFUs | *S. epidermidis* CFUs | *C. acnes*/ *S. epidermidis* CFU ratio | *ctm*+ / total *C. acnes* isolates | ppa0859 colony PCR results |
|---|---|---|---|---|---|---|
| 1a | 12 | 0 | 5 | 2.00 | 0/10 | Negative |
| 1b | 13600 | 0 | 10 | 1236.36 | 0/10 | Negative |
| 1c | 1880 | 0 | 112 | 16.64 | 0/10 | Negative |
| 1d | 49600 | 0 | 2 | 16533.33 | 0/10 | Negative |
| 1e | 1720 | 0 | 0 | 1720.00 | 0/10 | Negative |
| 1f | 245 | 0 | 0 | 245.00 | 0/10 | Negative |
| 1g | 4000 | 0 | 0 | 4000.00 | 0/10 | Negative |
| 1h | 110 | 51 | 0 | 110.00 | 1/10 | **Positive** |
| 1i | 10000 | 0 | 32 | 303.03 | 0/10 | Negative |
| 1j | 96000 | 0 | 6 | 13714.29 | 0/10 | Negative |
| 2a | 128000 | 0 | 0 | 128000.00 | 0/10 | Negative |
| 2b | 48000 | 0 | 0 | 48000.00 | 0/10 | Negative |
| 2c | 112000 | 0 | 1 | 56000.00 | 0/10 | Negative |
| 2d | 4000 | 0 | 2 | 1333.33 | 0/10 | Negative |
| 2e | 32000 | 0 | 0 | 32000.00 | 7/10 | **Positive** |
| 2f | 14000 | 0 | 1 | 7000.00 | 0/10 | Negative |
| 2g | 10800 | 0 | 0 | 10800.00 | 0/10 | Negative |
| 2h | 1200 | 0 | 11 | 100.00 | 0/10 | Negative |
| 2i | 1136000 | 0 | 0 | 1136000.00 | 10/10 | **Positive** |
| 2j | 29600 | 0 | 1 | 14800.00 | 0/10 | Negative |
| 3a | 4000 | 0 | 0 | 4000.00 | 0/10 | Negative |
| 3b | 184000 | 0 | 0 | 184000.00 | 0/10 | Negative |
| 3c | 5600 | 0 | 0 | 5600.00 | 0/10 | Negative |
| 3d | 10800 | 0 | 2 | 3600.00 | 0/10 | Negative |
| 3e | 56000 | 0 | 0 | 56000.00 | 0/10 | Negative |
| 3f | 104000 | 0 | 0 | 104000.00 | 0/10 | Negative |
| 3g | 312000 | 0 | 0 | 312000.00 | 0/10 | Negative |
| 3h | 200000 | 0 | 9 | 20000.00 | 0/10 | Negative |
| 3i | 1680 | 0 | 0 | 1680.00 | 0/10 | Negative |
| 3j | 44800 | 0 | 0 | 44800.00 | 0/10 | Negative |
| 4a | 8000 | 0 | 10 | 727.27 | 0/10 | Negative |
| 4b | 45000 | 0 | 0 | 45000.00 | 0/10 | Negative |
| 4c | 60000 | 5 | 0 | 60000.00 | 0/10 | Negative |
| 4d | 120000 | 0 | 14 | 8000.00 | 0/10 | Negative |
| 4e | 1000 | 0 | 0 | 1000.00 | 0/10 | Negative |
| 4f | 24000 | 0 | 50 | 470.59 | 0/10 | Negative |
| 4g | 320000 | 0 | 73 | 4324.32 | 0/10 | Negative |
| 4h | 88000 | 0 | 1 | 44000.00 | 0/10 | Negative |

| | | | | | | |
|---|---|---|---|---|---|---|
| 4i | 104000 | 0 | 65 | 1575.76 | 0/10 | Negative |
| 4j | 48000 | 0 | 10 | 4363.64 | 0/10 | Negative |
| 5a | 4800 | 0 | 2 | 1600.00 | 0/10 | Negative |
| 5b | 1300 | 0 | 1 | 650.00 | 0/10 | Negative |
| 5c | 14400 | 340 | 2 | 4800.00 | 10/10 | **Positive** |
| 5d | 580 | 0 | 2 | 193.33 | pooled | Negative |
| 5e | 3200 | 0 | 6 | 457.14 | pooled | Negative |
| 5f | 69600 | 0 | 10 | 6327.27 | pooled | Negative |
| 5g | 10000 | 0 | 0 | 10000.00 | pooled | Negative |
| 5h | 1200 | 0 | 0 | 1200.00 | pooled | Negative |
| 5i | 234000 | 0 | 6 | 33428.57 | pooled | **Positive** |
| 5j | 26000 | 0 | 2 | 8666.67 | pooled | Negative |
| 6a | 184000 | 0 | 0 | 184000.00 | 0/10 | Negative |
| 6b | 197000 | 8 | 18 | 10368.42 | 0/10 | Negative |
| 6c | 800000 | 0 | 0 | 800000.00 | 0/10 | Negative |
| 6d | 164000 | 1 | 1 | 82000.00 | 8/8 | **Positive** |
| 6e | 330000 | 1 | 1 | 165000.00 | 0/10 | Negative |
| 6f | 32000 | 14 | 40 | 780.49 | pooled | **Positive** |
| 6g | 92000 | 0 | 42 | 2139.54 | pooled | Negative |
| 6h | 168000 | 720 | 0 | 168000.00 | pooled | **Positive** |
| 6i | 196000 | 2 | 0 | 196000.00 | pooled | **Positive** |
| 6j | 224800 | 2 | 16 | 13223.53 | pooled | **Positive** |
| 7a | 880 | 0 | 0 | 880.00 | pooled | Negative |
| 7b | 16400 | 0 | 20 | 780.95 | pooled | Negative |
| 7c | 15800 | 0 | 1200 | 13.16 | pooled | Negative |
| 7d | 320 | 0 | 800 | 0.40 | pooled | Negative |
| 7e | 48800 | 0 | 0 | 48800.00 | pooled | Negative |
| 7f | 7000 | 0 | 2400 | 2.92 | pooled | Negative |
| 7g | 740 | 20 | 460 | 1.61 | pooled | Negative |
| 7h | 1600 | 0 | 1 | 800.00 | pooled | Negative |
| 7i | 4200 | 8 | 220 | 19.00 | pooled | Negative |
| 7j | 3200 | 40 | 0 | 3200.00 | pooled | Negative |
| 8a | 200 | 4600 | 100 | 1.98 | 0/10 | Negative |
| 8b | 5600 | 0 | 19 | 280.00 | 0/10 | Negative |
| 8c | 54 | 340 | 22 | 2.35 | 0/10 | Negative |
| 8d | 312000 | 0 | 300 | 1036.55 | pooled | Negative |
| 8e | 152000 | 0 | 100 | 1504.95 | pooled | Negative |
| 8f | 14800 | 160 | 200 | 73.63 | pooled | Negative |
| 8g | 334400 | 0 | 1800 | 185.67 | pooled | **Positive** |
| 8h | 70000 | 0 | 28 | 2413.79 | pooled | Negative |
| 8i | 24800 | 0 | 400 | 61.85 | pooled | Negative |

| | | | | | | |
|---|---|---|---|---|---|---|
| 8j | 768000 | 0 | 540 | 1419.59 | pooled | Negative |
| 9a | 128000 | 0 | 24 | 5120.00 | 0/10 | Negative |
| 9b | 64000 | 0 | 26 | 2370.37 | 0/10 | Negative |
| 9c | 100000 | 0 | 0 | 100000.00 | 0/10 | Negative |
| 9d | 2200 | 0 | 0 | 2200.00 | 0/10 | Negative |
| 9e | 112000 | 0 | 0 | 112000.00 | 10/10 | **Positive** |
| 9f | 6800 | 0 | 17 | 377.78 | 0/10 | Negative |
| 9g | 800000 | 0 | 0 | 800000.00 | 10/10 | **Positive** |
| 9h | 640000 | 0 | 12800 | 50.00 | 10/10 | **Positive** |
| 9i | 424000 | 0 | 0 | 424000.00 | 10/10 | **Positive** |
| 9j | 4000 | 0 | 920 | 4.34 | 2/10 | **Positive** |
| 10a | 7600 | 0 | 1700 | 4.47 | 0/10 | Negative |
| 10b | 550 | 0 | 410 | 1.34 | 0/10 | Negative |
| 10c | 310 | 0 | 0 | 310.00 | 0/10 | Negative |
| 10d | 4500 | 0 | 0 | 4500.00 | 0/10 | Negative |
| 10e | 840 | 0 | 7 | 105.00 | 0/10 | Negative |
| 10f | 28600 | 0 | 0 | 28600.00 | pooled | **Positive** |
| 10g | 7800 | 0 | 0 | 7800.00 | pooled | Negative |
| 10h | 5600 | 0 | 0 | 5600.00 | pooled | Negative |
| 10i | 3000 | 0 | 0 | 3000.00 | pooled | Negative |
| 10j | 400 | 0 | 0 | 400.00 | pooled | Negative |
| 11a | 1100000 | 0 | 0 | 1100000.00 | 0/10 | Negative |
| 11b | 90000 | 0 | 0 | 90000.00 | 0/10 | Negative |
| 11c | 106000 | 0 | 0 | 106000.00 | 0/10 | Negative |
| 11d | 120 | 0 | 0 | 120.00 | 0/6 | Negative |
| 11e | 180 | 0 | 0 | 180.00 | 0/9 | Negative |
| 11f | 262000 | 0 | 0 | 262000.00 | 0/10 | Negative |
| 11g | 140000 | 0 | 0 | 140000.00 | 0/10 | Negative |
| 11h | 176000 | 4400 | 0 | 176000.00 | 0/10 | Negative |
| 11i | 7400 | 200 | 0 | 7400.00 | 0/10 | Negative |
| 11j | 284000 | 340 | 0 | 284000.00 | 0/10 | Negative |
| 12a | 6615385 | 0 | 0 | 6615385.00 | 10/10 | **Positive** |
| 12b | 100000 | 0 | 0 | 100000.00 | 10/10 | **Positive** |
| 12c | 1138462 | 0 | 0 | 1138462.00 | 10/10 | **Positive** |
| 12d | 12307692 | 0 | 0 | 12307692.00 | 10/10 | **Positive** |
| 12e | 92 | 0 | 0 | 92.00 | 5/5 | **Positive** |
| 12f | 2508 | 0 | 0 | 2508.00 | 10/10 | **Positive** |
| 13a | 45077 | 1754 | 400 | 112.41 | 0/10 | Negative |
| 13b | 1461538 | 40000 | 62 | 23199.02 | 0/10 | Negative |
| 13c | 7385 | 0 | 31 | 230.78 | 0/10 | Negative |
| 13d | 18462 | 92 | 92 | 198.52 | 0/10 | Negative |

| | | | | | | |
|---|---|---|---|---|---|---|
| 13e | 11538 | 5831 | 138 | 83.01 | 0/10 | Negative |
| 13f | 14307692 | 0 | 46 | 304418.98 | 0/10 | Negative |
| 13g | 147692 | 15 | 462 | 318.99 | 0/10 | Negative |
| 13h | 11231 | 0 | 77 | 143.99 | 0/10 | Negative |
| 13i | 2415385 | 0 | 0 | 2415385.00 | 0/10 | Negative |
| 13j | 16769 | 708 | 31 | 524.03 | 0/10 | Negative |
| 14a | 56000 | 0 | 0 | 56000.00 | 1/10 | **Positive** |
| 14b | 1000000 | 40 | 0 | 1000000.00 | 0/10 | Negative |
| 14c | 1520000 | 0 | 0 | 1520000.00 | 0/10 | Negative |
| 14d | 820000 | 240000 | 0 | 820000.00 | 0/10 | Negative |
| 14e | 1000000 | 176000 | 0 | 1000000.00 | 0/10 | Negative |
| 14f | 2740000 | 0 | 0 | 2740000.00 | 0/10 | Negative |
| 14g | 2860000 | 0 | 0 | 2860000.00 | 0/10 | Negative |
| 14h | 16000000 | 0 | 0 | 16000000.00 | 0/10 | Negative |
| 14i | 246000 | 0 | 0 | 246000.00 | 0/10 | Negative |
| 14j | 1520000 | 200 | 0 | 1520000.00 | 0/10 | Negative |
| 15a | 11000000 | 20 | 0 | 11000000.00 | 10/10 | **Positive** |
| 15b | 42000000 | 60 | 380 | 110236.22 | 10/10 | **Positive** |
| 15c | 2180000 | 0 | 6200 | 351.56 | 0/10 | Negative |
| 15d | 1220000 | 20 | 0 | 1220000.00 | 0/10 | Negative |
| 15e | 1140000 | 0 | 84000 | 13.57 | 10/10 | **Positive** |
| 15f | 3640000 | 0 | 0 | 3640000.00 | 10/10 | **Positive** |
| 15g | 218000 | 0 | 3800 | 57.35 | 0/10 | Negative |
| 15h | 244000 | 0 | 0 | 244000.00 | 10/10 | **Positive** |
| 15i | 58000 | 0 | 0 | 58000.00 | 10/10 | **Positive** |
| 15j | 254000 | 0 | 0 | 254000.00 | 10/10 | **Positive** |
| 16a | 9800000 | 1760000 | 60 | 160655.74 | 0/10 | Negative |
| 16b | 20200000 | 10200000 | 0 | 20200000.00 | 0/10 | Negative |
| 16c | 1880000 | 17600 | 80 | 23209.88 | 0/10 | Negative |
| 16d | 51000000 | 4200000 | 114000 | 447.36 | 0/10 | Negative |
| 16e | 14600000 | 520000 | 180 | 80662.98 | 0/10 | Negative |
| 16f | 25600000 | 174000 | 102000 | 250.98 | 0/10 | Negative |
| 16g | 13000000 | 480 | 2120 | 6129.18 | 0/10 | Negative |
| 16h | 32000000 | 0 | 23400 | 1367.46 | 0/10 | Negative |
| 16i | 476000 | 420 | 17800 | 26.74 | 0/10 | Negative |
| 16j | 1250000 | 0 | 30200 | 41.39 | 0/10 | Negative |

**Table S6.** Cutimycin BGC presence/absence; the CFUs of *C. acnes*, *C. granulosum* and *S. epidermidis*; and the *C. acnes/S. epidermidis* CFU ratio in the content of individual human skin follicles.

| Bacterium or plasmid | Strain | Internal Reference | Characteristics | Source or Reference |
|---|---|---|---|---|
| **Plasmids** | | | | |
| pCGL0243 | | | *E. coli*- *C. glutamicum* shuttle vector | (*42*) |
| pJC215 | | | SuperCosI with the cutimycin BGC from *C. acnes* HL030PA1 | This study |
| pJS004 | | | pCGL0243 with the cutimycin BGC from *C. acnes* HL030PA1 | This study |
| **Species** | | | | |
| *Corynebacterium accolens* | ATCC 49725 | | | |
| *Corynebacterium glutamicum* | DSM 20300 | KPL1958 | Soil isolate derivative | |
| *Corynebacterium glutamicum* | | MF0704 | Heterologous host bearing pJS004 | This study |
| *Corynebacterium jeikeium* | DSM7171 | | | |
| *Corynebacterium striatum* | DSM20668 | | | |
| *Corynebacterium pseudodiphtheriticum* | DSM44287 | | | |
| *Cutibacterium acnes* | HL030PA1 | | | |
| *Cutibacterium acnes* | HL053PA2 | | | |
| *Cutibacterium acnes* | HL086PA1 | KPL2374 | | |
| *Cutibacterium acnes* | HL110PA1 | | | |
| *Cutibacterium acnes* | KPA171202 | KPL2017 | | |
| *Staphylococcus aureus* | JE2 | | | |
| *Staphylococcus aureus* | UAMS-1 | | | |
| *Staphylococcus aureus* | USA300 NRS384 | | | |
| *Staphylococcus epidermidis* | ATCC 35984 | | | |
| *Staphylococcus epidermidis* | DSM20042 | | | |
| *Staphylococcus epidermidis* | W23144 | | | |

**Table S7.** Bacterial strains and plasmids used in this study.

| **Primer name** | **5'-3' nucleotide sequence** |
|---|---|
| JC_Super_HL030_FWD | GTATTCGGAGGCTGGAAGACTATCGTCGCCGCACT |
| JC_Super_HL030_REV | GCGGTAGTTGTCGCACTATAGGGATCCTTTAGTGAGGGT |
| JC_HL030_pt1_FWD | ACTAAAGGATCCCTATAGTGCGACAACTACCGCCTACTCC |
| JC_HL030_pt1_REV | GCAGCCTCGTGGAAGAGATCCCGTAAATCGCGCCAAGTCG |
| JC_HL030_pt2_FWD | CGATTTACGGGATCTCTTCCACGAGGCTGCTACTGTTGC |
| JC_HL030_pt2_REV | GGCGACGATAGTCTTCCAGCCTCCGAATACATCTCAAC |
| oKL244 | GCAGAATAAATGATCCGTCGAG |
| oKL425 | ACTATCCGCAAGCGCGAA |
| oKL439 | TAGCGCTCTGGGGTGGGA |
| oKL535 (ppa0859_F) | ACCAGCAGGCTTACGGC |
| oKL536 (ppa0859_R) | GGTCACTGTGGAGCTGG |
| oKL564 (*Cac* gyrB_F) | GAGCATCGTCCGAAAGTCAC |
| oKL565 (*Cac* gyrB_R) | TCGCCCTCCACAATGAAGAT |
| oKL572 (ppa0860_F) | GTTGCATGCCATCTCCGTAG |
| oKL573 (ppa0860_R) | ACCGATCCGCCATACTTTCT |

**Table S8**. Primers used in this study.

| | Fragment | Calculated *m/z* | Observed *m/z* |
|---|---|---|---|
| 1 | b15 | 1113.3268 | 1113.32994 |
| 2 | b14 | 1044.3053 | 1044.30782 |
| 3 | b7-b10 | 912.2735 | 912.27479 |
| 4 | b6-b10 | 813.2045 | 813.20602 |
| 6 | b6-b7 | 1032.2689 | 1032.27121 |
| 7 | b3-b7 | 799.1889 | 799.1905 |
| 8 | b3-b6 | 898.2573 | 898.25602 |
| 9 | b3-b9 | 649.146 | 649.14485 |
| 10 | b3-b10 | 580.1245 | 580.12535 |
| 11 | b2-b9 | 483.1937 | 483.20071 |
| 12 | b3-b7 | 333.1557 | 333.15563 |
| 13 | b3-b6 | 234.0873 | 234.0866 |
| 14 | valine-imminium | 72.0808 | 72.08003 |

**Table S9.** The key fragments confirming the order of the amino acids in cutimycin.
